# KTD1 is a yeast defense factor against K28 killer toxin

**DOI:** 10.1101/2021.10.25.465803

**Authors:** Ilya Andreev, Simone M. Giovanetti, Guillaume Urtecho, Daniel Shriner, Joshua S. Bloom, Meru J. Sadhu

**Affiliations:** Genetic Disease Research Branch, National Human Genome Research Institute, National Institutes of Health, Bethesda, MD 20892, USA; Molecular Biology Interdepartmental Doctoral Program, University of California, Los Angeles, Los Angeles, CA 90095, USA; Center for Research on Genomics and Global Health, National Human Genome Research Institute, National Institutes of Health, Bethesda, MD 20892, USA; Department of Human Genetics, University of California, Los Angeles, Los Angeles, CA 90095, USA; Department of Biological Chemistry, University of California, Los Angeles, Los Angeles, CA 90095, USA; Howard Hughes Medical Institute, University of California, Los Angeles, Los Angeles, CA 90095, USA; Institute for Quantitative and Computational Biology, University of California, Los Angeles, Los Angeles, CA 90095, USA; Department of Computational Medicine, University of California, Los Angeles, Los Angeles, CA, 90095, USA

## Abstract

Secreted protein toxins are widely used weapons in conflicts between organisms. Killer yeast produce killer toxins that inhibit the growth of nearby sensitive yeast. We investigated variation in resistance to the killer toxin K28 across diverse natural isolates of the *Saccharomyces cerevisiae* population and discovered a novel defense factor, which we named *KTD1*, that is an important determinant of K28 toxin resistance. *KTD1* is a member of the DUP240 gene family of unknown function. We uncovered a putative role of DUP240 proteins in killer toxin defense and identified a region that is undergoing rapid evolution and is critical to *KTD1*’s protective ability. Our findings implicate *KTD1* as a key factor in the defense against killer toxin K28.

## Introduction

Conflict between organisms is a major evolutionary force, both within and between species, and can lead to complex molecular arms races of competing evolutionary adaptation. Secreted protein toxins are a common weapon in biological conflict, encountered across all domains of life (*1–3*). Exposure to these toxins can select for adaptations in targeted organisms that allow them to resist toxic effects (*4–11*).

Toxin-secreting killer yeast strains of *Saccharomyces cerevisiae* are a well-studied example of conflict in the microbial world (*12*). They are infected by dsRNA killer viruses, which encode killer toxins that the killer yeast secrete to inhibit the growth of nearby susceptible yeast strains. As killer viruses also protect their hosts from the secreted toxins, a killer virus infection is beneficial to the host in competition with sensitive cells (*13–15*). The killer toxin K28, encoded by the M28 killer virus, enters target cells by hijacking their retrograde trafficking pathway, ultimately translocating to the cytoplasm and inducing G1/S cell cycle arrest through an unknown mechanism (*16*). The biology of K28 toxicity has been extensively studied; however, it is unknown whether natural populations of *S. cerevisiae* have adapted strategies of resistance to this killer toxin, if at all. Here, we leveraged natural variation in yeast to investigate whether cells have evolved mechanisms of protection against K28.

## Results

We surveyed K28 killer toxin resistance in a diverse panel of 16 *S. cerevisiae* strains isolated from a variety of geographical locations and ecological settings (*17*) (Fig. 1A; table S1) and found that they ranged from completely resistant to highly sensitive to K28 (Fig. 1B, fig. S1), consistent with past reports of variation in killer toxin resistance (*18–20*). Yeast infected with M28 killer virus are resistant to K28, as killer viruses confer toxin self-immunity upon the host cell (*21*). However, we did not detect M28 killer virus in any of these strains (figs. S2, S3), in keeping with previous findings that most yeast isolates are not infected by killer viruses (*19, 20, 22, 23*). This suggests that differences in toxin resistance are due to variation in chromosomally encoded genetic factors.

**Figure 1.**
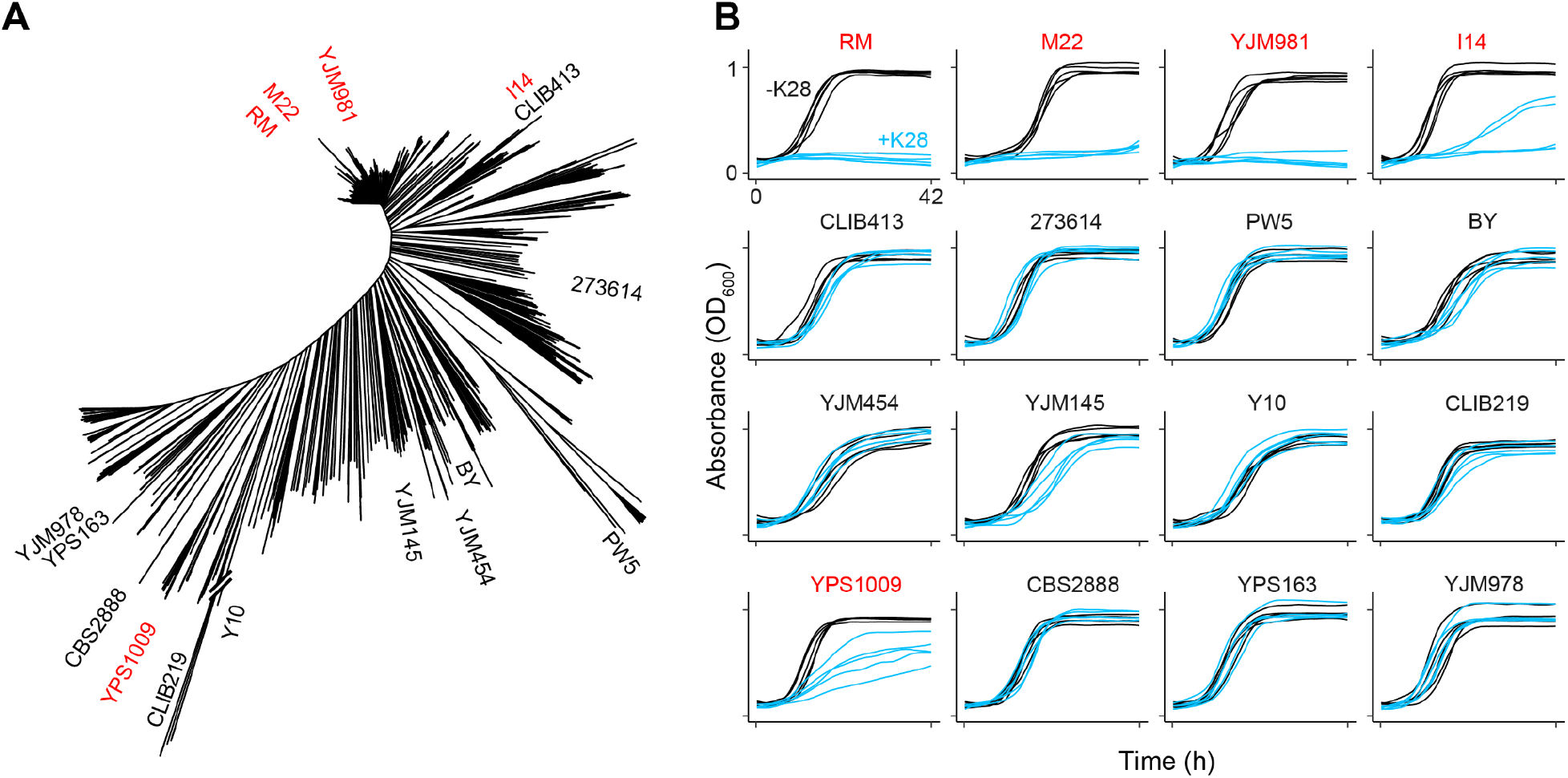
High natural variation in K28 resistance. **(A)** Neighbor-joining phylogenetic tree of 1,011 budding yeast isolates (*30*), highlighting the 16 strains selected for our panel. Name coloring reflects K28 killer toxin resistance, with red labels for sensitive strains and black labels for resistant strains, as determined in part B. **(B)** Growth curves of 16 diverse *S. cerevisiae* strains in minimal media with or without K28 (blue and black, respectively; *n* = 5). K28 resistance was quantified as the ratio of the areas under the growth curves in media containing or lacking K28 (AUC_+K28_ / AUC_-K28_) (fig. S1). Statistical analysis was performed using one-way analysis of variance (ANOVA) followed by Tukey’s post-hoc HSD test, and identified five strains as K28-sensitive (strain names in red) in comparisons with BY (*P* adjusted < 6.6×10^-6^).

To identify genes controlling variation in toxin resistance, we initially focused on the K28-resistant lab isolate BY and the highly sensitive wine isolate RM (table S1). We determined the toxin resistance of 912 fully genotyped haploid segregants generated from a BY × RM cross (*24*) by comparing growth in media with or without K28 toxin (Fig. 2A). Linkage analysis revealed three quantitative trait loci (QTLs) associated with K28 resistance (Fig. 2B, table S2). The strongest QTL was on chromosome I: only 2% of segregants with the RM allele grew robustly in the presence of K28, as opposed to 40% of the segregants with the BY allele (Fig. 2C). The 95% confidence interval for this QTL was 4.0 kb, with the peak located between the genes *UIP3* and YAR028W. Both genes are members of the DUP240 gene family of unknown function (*25, 26*).

**Figure 2.**
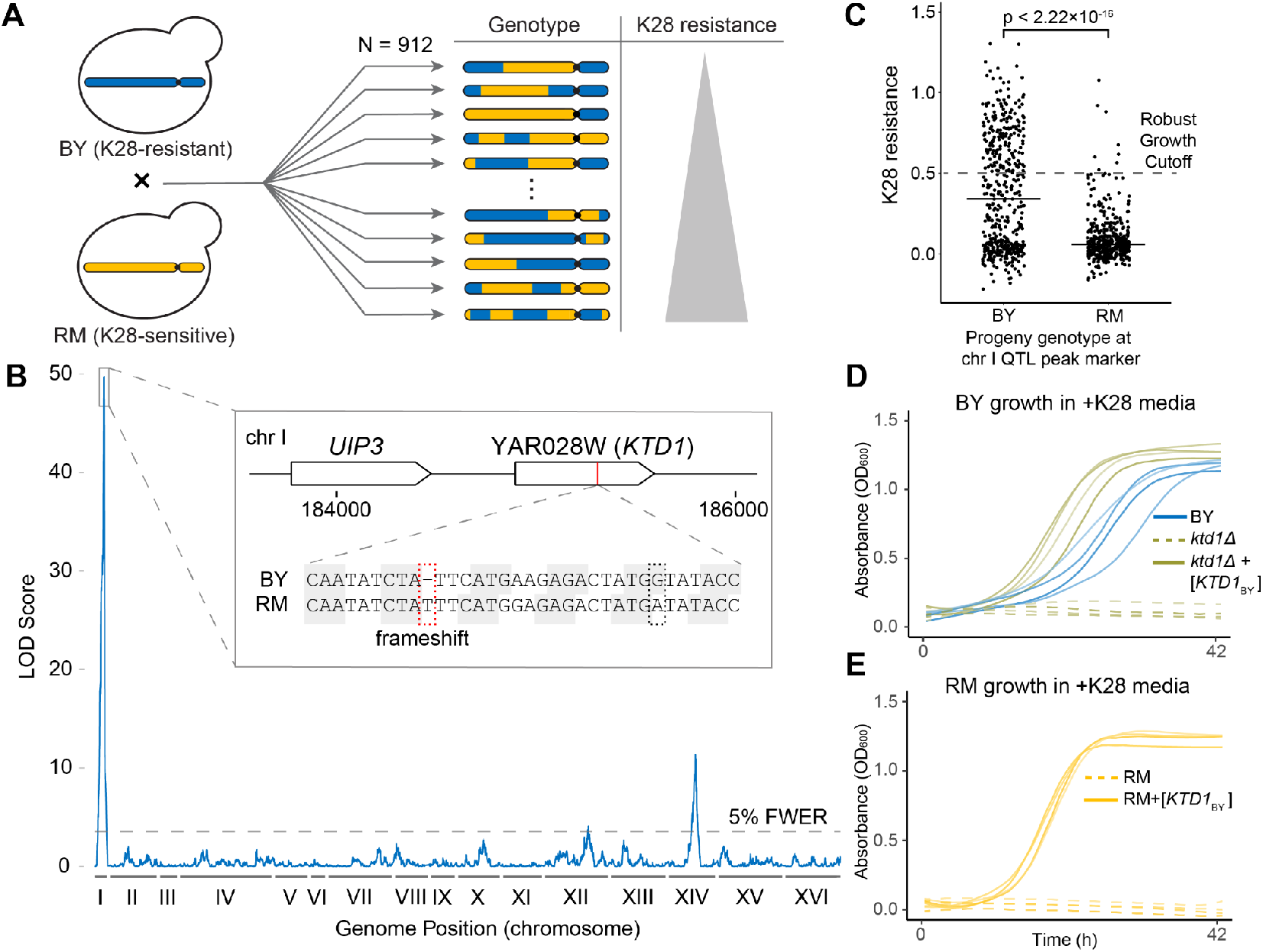
QTL mapping reveals YAR028W (*KTD1*) as a major genetic factor in killer toxin resistance. **(A)** Schematic representation of the workflow leading up to QTL mapping. 912 recombinant haploid progeny from a cross between BY and RM were assayed for K28 resistance, measured as AUC_+K28_ / AUC_-K28_. **(B)** QTL mapping results. Shown is the degree of association of a genomic locus with K28 resistance phenotype, measured as the LOD (logarithm of the odds) score, plotted against genomic coordinates. Three significant QTLs were found above the 5% family-wise error rate (FWER) significance threshold (LOD = 3.55, gray dashed line). The inset shows the genomic context of the QTL peak on chromosome I. **(C)** Phenotypic distribution of 912 BY × RM progeny grouped by their genotype at the peak marker for the chr I QTL. *P* < 2.22×10^-16^ by Welch’s two-sample *t*-test. The solid horizontal black lines denote mean phenotypes. We used an AUC_+K28_/AUC_-K28_ cutoff of 0.5 to define robust growth. **(D)** Growth of BY with an empty vector, and of BY *yar028wΔ* (*ktd1Δ*) carrying either an empty vector or YAR028W_BY_. Strains were grown in K28-containing media. Four biological replicates are shown in each plot. **(E)** Growth of RM carrying either an empty vector or the YAR028W_BY_ (*KTD1*_BY_) allele. In D and E, strains with YAR028W_BY_ (*KTD1*_BY_) were significantly more K28-resistant than those without, *P* < 0.005 by Welch’s two-sample one-tailed *t*-test (fig. S6).

We examined *UIP3* and YAR028W. RM’s YAR028W allele, YAR028W_RM_, had a 1-nucleotide frameshift mutation at codon 136 (Fig. 2B, inset), leading to 13 additional out-of-frame codons and a premature stop; in comparison, BY’s intact allele is 235 codons long. In contrast, both strains had full-length *UIP3* alleles. Analysis of ribosome footprinting data collected by Albert, et al. (*27*) showed that YAR028W is expressed and translated in BY, but its translation ends prematurely in RM, near the frameshift mutation (fig. S4). Furthermore, a genome-wide screen of 4806 deletion mutants in the yeast knockout collection, which is isogenic to BY, annotated the YAR028W deletion mutant, *yar028wΔ,* as partially sensitive to K28 (*28*). We confirmed that *yar028wΔ* is sensitive to K28, whereas we observed no effect of deleting *UIP3* (Fig. 2D and figs. S5, S6). Importantly, plasmid-based expression of the BY allele of YAR028W, YAR028W_BY_, in the sensitive RM and *yar028wΔ* strains provided strong protection against K28 (Fig. 2, D and E, fig. S6), while the RM allele had no effect (fig. S6). We concluded that YAR028W underlies the dramatic difference in K28 resistance between BY and RM. Therefore, we designated YAR028W as *KTD1* (*k*iller *t*oxin *d*efense).

We examined the molecular defense mechanism of Ktd1p. The first step in the process of K28 cytotoxicity is K28 binding to the cell wall (*16*), so we tested whether Ktd1p counters K28 at this step or after. Using a toxin adsorption assay (*28, 29*), we assayed the ability of K28 to bind to the surface of strains with or without *KTD1*_BY_ and in different genetic backgrounds. We found that expression of *KTD1*_BY_ did not reduce K28 binding (fig. S7). Additionally, toxin-sensitive RM did not exhibit higher K28 binding than toxin-resistant BY. As a control, we recapitulated that deletion of the mannosyltransferase *MNN2* prevents K28 binding (*28*). We therefore concluded that *KTD1* functions after the initial step of toxin binding to the cell surface.

We next explored the connection between variation in *KTD1* and K28 resistance across the yeast population. We found that 37% of 1,011 *S. cerevisiae* strains sequenced in a recent population survey had a premature stop mutation at codon 141 (*30*) (Fig. 3A). This polymorphism is also present in *KTD1*_RM_ (Fig. 2B, inset, black dashed box), indicating it was already truncated prior to acquiring the frameshift mutation. Examining the strains in our panel, the strain YJM981 carried the *ktd1-*141Stop mutation without the frameshift (fig. S8) and was also highly sensitive to K28. Two other sensitive strains in our panel, YPS1009 and I14, had full-length *KTD1* alleles, albeit with multiple nonsynonymous mutations in each (fig. S8). We tested whether any of these *KTD1* alleles conferred toxin resistance upon the sensitive *ktd1Δ* strain. Expression of *KTD1*_YJM981_ or *KTD1*_YPS1009_ did not protect against K28 toxin (Fig. 3B). However, *KTD1*_I14_ did confer K28 resistance, indicating that strain I14’s toxin sensitivity is caused by a different mechanism despite carrying a protective *KTD1* allele. We tested three additional *KTD1* alleles shared between five K28-resistant strains and found that all of them restored toxin resistance in the *ktd1Δ* strain (Fig. 3B). Our results indicate that *KTD1* function is a major determinant of variation in K28 resistance across the yeast population.

**Figure 3.**
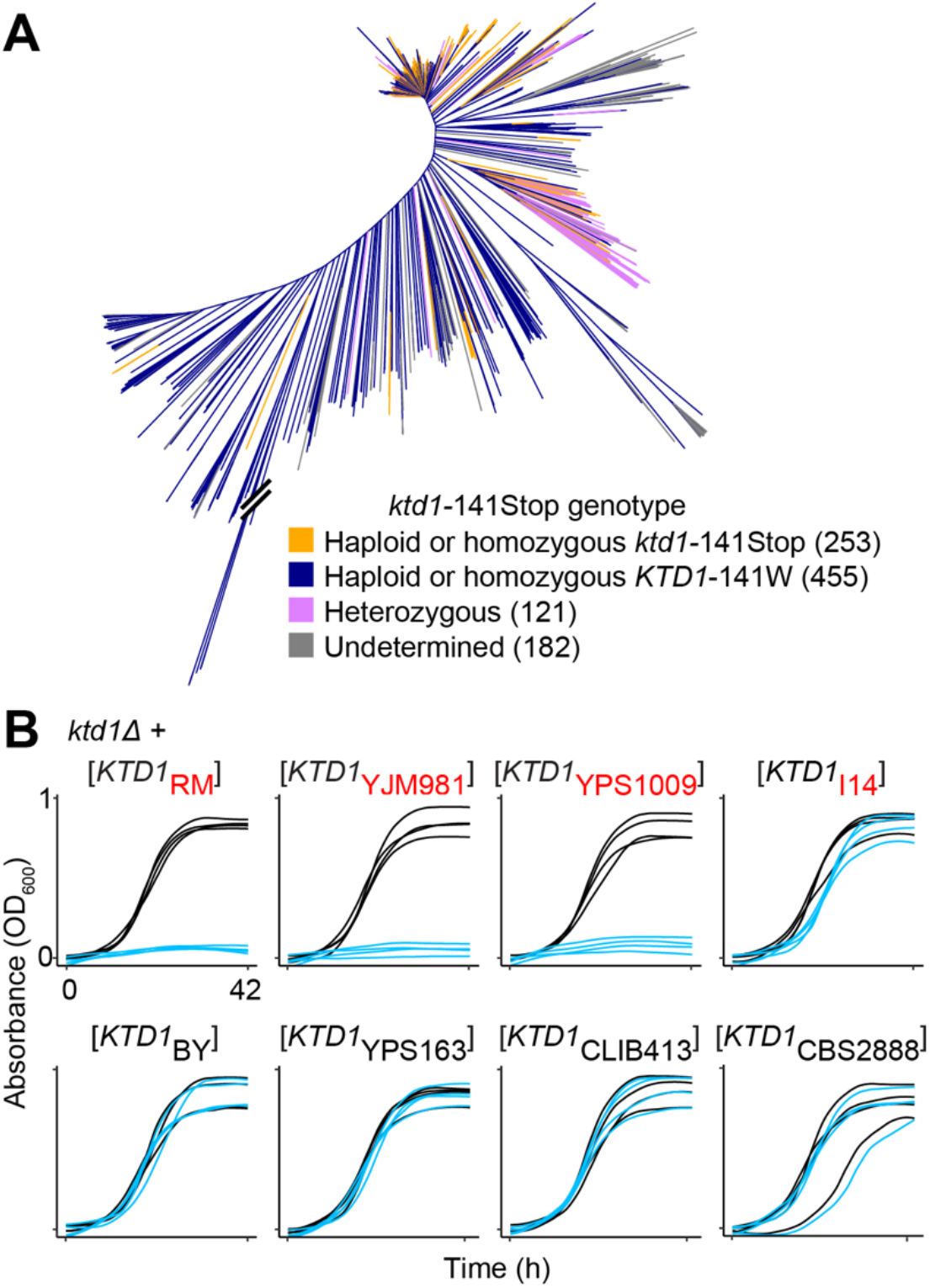
Analysis of *KTD1* alleles in the yeast population. **(A)** Distribution of the *ktd1*-141Stop mutation among 1,011 yeast isolates (*30*). **(B)** Growth curves of strains carrying the indicated *KTD1* allele in the BY *ktd1Δ* background; blue and black growth curves correspond to strains grown in minimal media with or without K28, respectively (*n* = 4). The subscript of the allele label indicates the strain in which that *KTD1* allele was found, while color indicates that strain’s phenotype (black, resistant; red, sensitive). ANOVA followed by Tukey’s HSD (fig. S8) identified the first three alleles as significantly less resistant than the last five (*P* adjusted < 5×10^- 14^ for all between-group comparisons).

*KTD1* is a member of the enigmatic DUP240 gene family. None of the 10 DUP240 genes in the reference genome have a clearly ascribed function (table S3), and simultaneous deletion of all DUP240 genes does not affect yeast growth (*25*). Moreover, this family recently arose in the *Saccharomycetaceae* family (fig. S9), and the number and composition of this family varies widely between strains (*26*) (table S4). DUP240 proteins have two predicted transmembrane domains separated by a short inter-helix linker. To identify the protein domains responsible for protection, we constructed chimeras between Ktd1p and Uip3p (Fig. 4A), the latter of which is the closest DUP240 family member with 58.1% amino acid identity to Ktd1p (fig. S10) but does not confer K28 resistance. Our experiments identified two regions of Ktd1p important for protection: the inter-helix linker and the C-terminus (Fig. 4A). When Ktd1p’s 88 C-terminal amino acids were replaced with the corresponding region of Uip3p (chimera K/U-2), K28 resistance was fully abrogated. In addition, chimeras U/K-5 and U/K-6, which differed only in the identity of their inter-helix linker, showed very different K28 protection responses (Fig. 4A), indicating a key role for Ktd1p’s inter-helix linker in K28 protection. The inter-helix linker region was of particular interest, as it is predicted to reside on the non-cytoplasmic face of the membrane (*31*) and thus is the only part of Ktd1p that could reside in the same cellular compartment as K28 before it reaches the cytoplasm (*16*).

**Figure 4.**
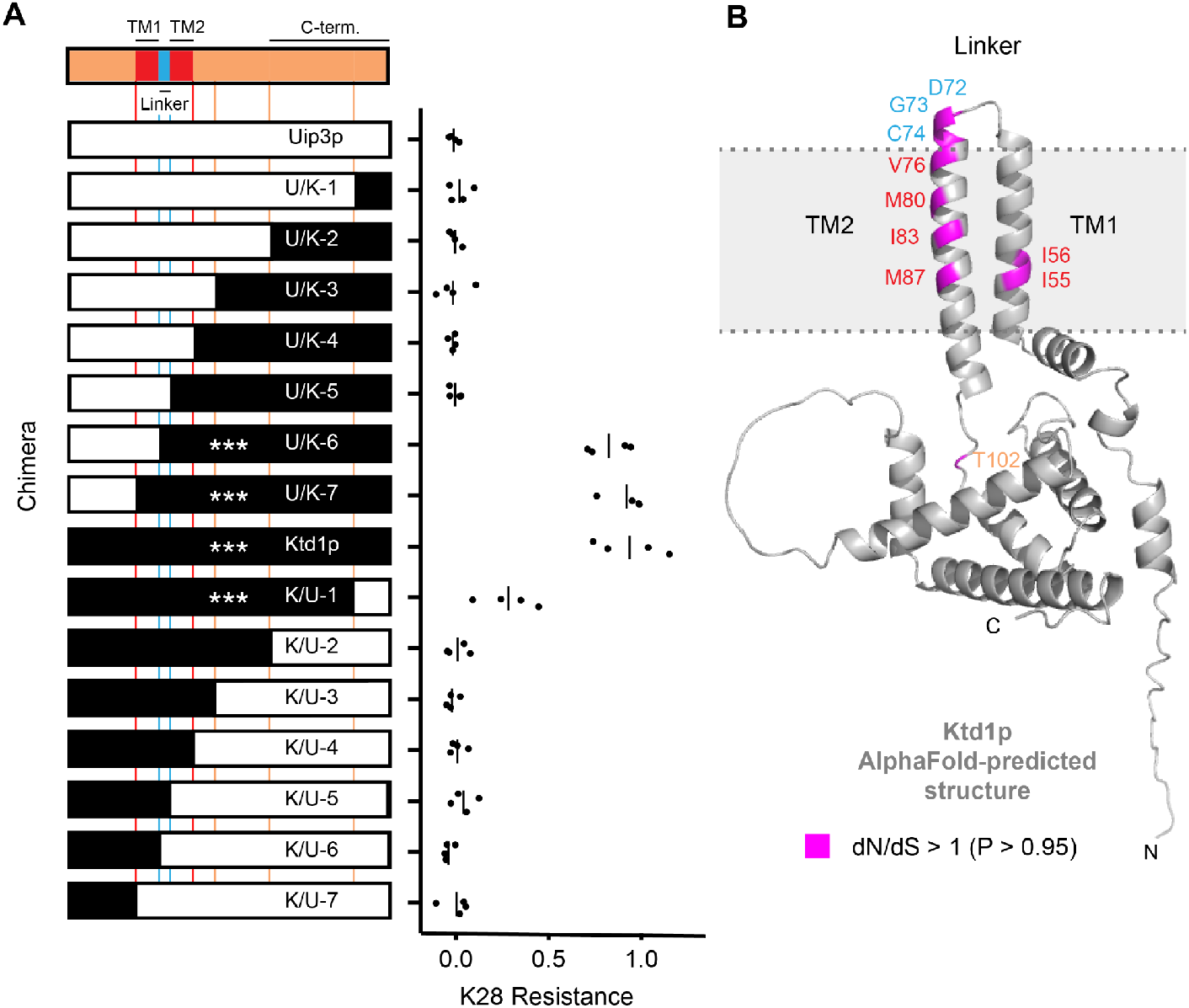
Ktd1p protein analysis. **(A)** K28 resistance phenotypes of chimeras between Ktd1p and Uip3p expressed in the BY *ktd1Δ* background. White segments denote regions of chimeras derived from the non-protective Uip3p, and black segments denote regions derived from the protective Ktd1p. Four biological replicates are shown for each chimera’s K28 resistance phenotype, measured as AUC_+K28_ / AUC_-K28_ from the growth curves in fig. S10, with vertical bars denoting the sample mean. \*\*\**P* adjusted < 0.001 in comparison to Uip3p (ANOVA followed by Tukey’s HSD). **(B)** Sites of positive selection in Ktd1p. Sites having a greater than 0.95 probability of dN/dS above 1 are highlighted in magenta. Protein structure of Ktd1p was predicted by AlphaFold (*37*). The colors of the amino acid labels correspond to the coloring of the topological domains of Ktd1p in part A, as defined by Poirey, et al. (*25*).

We noted that the linker and its surrounding sequence are highly diverged between DUP240 family members (fig. S11). Proteins involved in host defense often experience evolutionary pressure to change (*32*), which at the genetic level can manifest as a higher nonsynonymous mutation rate relative to synonymous mutation rate (dN/dS). We searched the genomes of our 16-isolate yeast panel for ORFs encoding proteins homologous to known DUP240s and selected 18 orthologs to generate models of dN/dS ratios per codon using the codeml package in PAML (*33*). Ten codons were identified as having dN/dS > 1, indicating positive selection at those sites (table S5). The corresponding residues are concentrated in the region spanning the transmembrane helices and the linker (Fig. 4B, fig. S11). Combined, the evidence of positive selection in the Ktd1p inter-helix linker and its role in K28 protection suggest that the DUP240 family evolved under selective pressure from killer toxins.

## Discussion

Protein toxins are powerful weapons in inter- and intra-species competition that can lead to polymorphic adaptations in the targeted populations (*4–7, 10, 34*). We discovered a polymorphic yeast gene, *KTD1*, that protects yeast against the killer toxin K28. *KTD1* is a member of the enigmatic DUP240 gene family, a rapidly expanding and contracting family of genes with unknown function. We found many distinct alleles of *KTD1* in the yeast population, including a highly prevalent premature stop codon, which could reflect coevolution with K28, varying across the diverse environments and geographical locations in which yeast are found. *KTD1* is under positive selection at the molecular level, concentrated in amino acids in the transmembrane domains and the linker between them, which is critical for Ktd1p-mediated resistance to K28. After binding to the cell surface, K28 is endocytosed and then avoids vacuolar degradation (*16*), instead travelling to the ER before entering the cytoplasm. Ktd1p might provide resistance by facilitating vacuolar degradation of K28, consistent with our finding that Ktd1p acts after K28 binds to the cell surface and the involvement of other DUP240s in protein trafficking (*35*) and the related DUP380s in mediating degradation of endocytosed proteins (*36*).

## Acknowledgments

We thank Keith Kozminski, Leonid Kruglyak, Peter Stirling, Frank Albert, Helen Murphy, and members of the Sadhu and Kruglyak labs for helpful discussion. We thank David Drubin, Leonid Kruglyak, and Reed Wickner for strains and plasmids.

## Funding

National Institutes of Health grant 1ZIAHG200401 (IA, SMG, MJS)

National Institutes of Health grant 5T32GM007185 (GU)

National Institutes of Health grant 1ZIAHG200362 (DS)

Howard Hughes Medical Institute (JSB)

## Author contributions

Conceptualization: MJS

Methodology: IA, SMG, GU, DS, JSB, MJS

Investigation: IA

Formal analysis: IA

Visualization: IA, MJS

Supervision: MJS

Resources: JSB

Writing – original draft: IA, MJS

Writing – reviewing & editing: IA, SMG, MJS

## Data and materials availability

All data and code will be archived in a permanent, independent repository prior to publication. Currently, all data and code are located at https://github.com/ilya-andreev2/killer-virus.

## Materials and Methods

### Strains

Strains, plasmids, and oligonucleotide sequences used in this study are listed in Tables S6, S7, and S8, respectively. All constructed plasmids were sequence-verified using Sanger sequencing (Eurofins Genomics, Louisville, KY, USA).

The 16-isolate panel of diverse *S. cerevisiae* strains, MSY24-MSY39, was composed of stable haploid strains made by Bloom, et al. (*17*) and were gifts from Leonid Kruglyak, as well as MSY1 and MSY8 and the panel of BY × RM segregants (*24*). MS300c (MSY20) and 192.2d (MSY21) were gifts from David Drubin (*28*).

We generated a diploid K28-secreting strain for production of toxin-containing supernatant (described in the section “Preparation of Toxic and Non-toxic Supernatants” below). As this supernatant was to be applied to haploid cells of either a or alpha mating type, we designed our K28-secreting strain to be diploid so that it would not release mating pheromone into the media. A K28 hypersecretor, MS300c (MATalpha *leu2 ski2-2* {M28 infected}) (*38*), was mated to *ski2Δ* (MATa *his3Δ leu2Δ met15Δ ura3Δ ski2Δ*::KanMX) from the MATa knockout collection (Transomic, Huntsville, AL) to produce a hypersecretor diploid *(ski2Δ*/*ski2-2*) strain, MSY52, that is infected with the M28 virus and preserves the hypersecreting phenotype due to the absence of *SKI2* function.

We generated a virus-cured strain for production of non-toxic control media. Virus curing was performed by growth of MSY52 at high temperature (*39*). MSY52 was pre-grown in liquid YPD media overnight at 37 C (elevated) or 30 C (control). Cultures were diluted in fresh YPD to a density of approximately 500 cells/200 μL, at which point 200 μL of cultures were pipetted onto YPD agar plates. After a 2-day incubation at respective temperatures (30 or 37 C), colonies were replica-plated onto Methylene Blue Agar (MBA) plates (pH 4.7) seeded with lawns of the K28-hypersensitive 192.2d strain (MSY21). All 30 C control colonies showed killer activity, indicated by the inability of 192.2d to grow in their vicinity. Several colonies from the 37 C plate no longer exhibited killer phenotype, one of which was selected and named MSY53. Absence of M28 virus was confirmed by virus typing assay (see below).

*uip3Δ*, *ktd1Δ* (MSY123), and *mnn2Δ* were taken from the MATa knockout collection.

The remaining strains in Table S6 were generated by transforming *ktd1Δ*, *mnn2Δ*, RM (MSY8), or BY (MSY1) with the plasmids described in Table S7 using standard lithium acetate transformation procedures (*40*).

### Stock Solutions and Media

Phosphate-Citrate Buffer (pH 4.7) was prepared by first dissolving 56.88 g K_2_HPO_4_ into 200 mL distilled water and subsequently titrating with approximately 30 g citric acid until pH 4.7 was reached. The solution was sterilized by autoclaving for 20 min. Crystals may form over long periods of storage, which can be redissolved by heating the buffer to 80 C for 20-30 min, stirring, and cooling.

Specialized low-pH CSM minimal media for production of K28 killer toxin, CSM pH 4.7 + 0.05% gelatin, was prepared by adding pre-mixed CSM powders (Sunrise Science, Knoxville, TN) and YNB (BD, Franklin Lakes, New Jersey, USA) according to manufacturer instructions, dextrose to 2% (MP Biomedicals, Irvine, CA), and adding back uracil and leucine to the concentrations specified in Dunham, et al. (*41*). pH was adjusted with 112 mL of pH 4.7 phosphate-citrate buffer per 1 L media. For additional killer toxin stability, media was supplemented with 0.05% type B gelatin (MilliporeSigma, Burlington, MA, USA, Cat. #G9391-100G) as per Woods and Bevan (*42*). The mixture was heat-stirred for 2 h to allow gelatin to dissolve. All CSM media were sterilized using 0.22 µm polyethersulfone membrane filters (MilliporeSigma).

For use with supernatants, as described below, we made concentrated replenishment minimal media, 5xCSM pH 4.7, either with or without histidine as appropriate for the experiment. These media were prepared by dissolving either 0.8375 g CSM-Leu-Ura or 0.8125 g CSM-His-Leu-Ura powders (Sunrise Biosciences) in 222 mL distilled H_2_O, along with 125 mg leucine, 25 mg uracil, 8.375 g YNB, and 25 g dextrose. pH was adjusted with 28 mL phosphate-citrate buffer (pH 4.7). The solutions were stirred for approximately 20 min to ensure full dissolution of uracil crystals and subsequently filter-sterilized.

YPD media contained 1% yeast extract (Thermo Fisher Scientific, Waltham, MA, USA), 2% peptone (BD), and 2% dextrose, and was sterilized by autoclaving. When used for plates, the media additionally contained 2% agar.

YPD-based Methylene Blue Agar (MBA) plates (pH 4.7) were prepared as described in Amberg et al. (*43*). Briefly, 1% yeast extract, 2% peptone, 2% glucose, and 2% agar were dissolved in 415 mL distilled water and autoclaved for 20 min. Following autoclaving, media was supplemented with sterile solutions of 25 mL 40% glucose, 4.2 mL of 4 mg/mL methylene blue dye (MilliporeSigma), and 56 mL phosphate-citrate buffer of pH 4.7. Correct pH was confirmed using a pH probe after buffer addition.

### Preparation of Toxic and Non-toxic Supernatants

Single colonies of MSY52 (diploid K28-secretor) or MSY53 (virus-free diploid) were separately inoculated into 200 mL of CSM pH 4.7 + 0.05% gelatin and incubated for 24-48 h at 23 C, 70 RPM. These conditions were necessary for generating stable, high-activity K28 killer toxin. For experiments with strains carrying a *HIS3* plasmid, CSM-His was used instead of CSM for supernatant preparations. Cultures were filtered with 0.22 µm polyethersulfone membrane filters into pre-chilled glass bottles, producing the toxic (MSY52) and non-toxic (MSY53) supernatants.

### Liquid Growth Assay for Killer Toxin Resistance

Single colonies of yeast strains were inoculated and pre-grown in 100 μL CSM or CSM-His (pH 7.0) for 24 h in CORNING 96-well plates prior to the experiment. 1.5 μL of saturated yeast cultures were transferred into 160 μL of either the toxic or non-toxic supernatant, supplemented with 40 μL 5xCSM pH 4.7 (with or without histidine) to replenish spent nutrients, for a total volume of 201.5 μL in each well. Strains were randomly distributed across the plate to avoid confounding of strain identity with position on plate. The one exception is the experiment comparing toxin sensitivity of *uip3Δ* and *ktd1Δ* to WT (fig. S5), in which each strain occupied a specific row on the plate. Absorbance (OD_600_) of each well was measured on a SPECTROstar Omega microplate reader equipped with a plate stacker (BMG Labtech, Ortenberg, Germany) over a period of 2-3 days at ambient temperature (∼25 C) with 30 s of shaking at 600 RPM prior to each plate reading. Reads were taken with variable frequency depending on the number of plates being read, with cycle time ranging from 8 to 104 minutes. For each strain in an individual well, the K28 toxin resistance phenotype was calculated, following blank correction, as the ratio of the area under the growth curve in toxic media (AUC_+K28_) to the area under the growth curve in non-toxic media (AUC_-K28_), i.e. AUC_+K28_/AUC_-K28_. Values close to 0 indicate low killer toxin resistance, while values close to 1 correspond to high resistance.

### Killer spotting assay

4 mL YPD cultures of MSY21, MSY52-53, and MSY24-39 were grown overnight, with MSY39 (ade^-^) supplemented with 1% adenine. 2×10^7^ or 4×10^7^ cells of K28-hypersensitive MSY21 (192.2d) were uniformly spread on 10-cm MBA agar plates, with 3 replicate plates for each plating density. Cultures of MSY52 (K28-secreting control), MSY53 (virus-free control), and MSY24-39 (the 16-isolate panel) were pelleted at 3000 rpm and resuspended in fresh YPD to make a ∼50% v/v cell slurry, as previously described (*28*). 150 μL of these slurries were pipetted into wells of a 96-well plate and pinned with a VP407AH metal replicator (V&P Scientific, San Diego, CA) to transfer approximately 3 μL of slurry onto the previously MSY21-seeded MBA plates. Each spot of pinned cells was imaged through microscope oculars using the back camera of a Galaxy S10e mobile phone (Samsung, Seoul, South Korea) with Open Camera v1.49.1 software, which allowed imaging of colonies at fixed exposure and white balance within an individual plate.

### Viral RNA typing assay

To assay for presence of M28 virus in a given strain, RT-PCR-based virus typing was performed as described by Chang, et al. (*18*). Briefly, following isolation of total RNA with the Qiagen RNeasy Mini Kit (Qiagen, Valencia, CA, USA), cDNA was synthesized by incubating RNA for 2 h at 37 C using the High-Capacity cDNA Reverse Transcription Kit (Applied Biosystems, Foster City, CA, USA). Afterwards, PCR with Kapa HiFi HotStart ReadyMix (Roche, Basel, Switzerland) was performed on the cDNA using M28 virus-specific primer pairs: oIA9/oIA10, oIA11/oIA12, and oIA13/oIA14 with expected fragment sizes of 253, 180, and 350 bp, respectively. These primers correspond exactly to M28-F1/R1, M28-F2/R2, and M28-F3/R3 primers used by Chang, et al. Presence of M28 virus was characterized by strong PCR bands at expected amplicon lengths. A band of low intensity at 650 bp appeared in several experiments, which we deemed an artifact because its presence was irrespective of the particular PCR primer pair.

### QTL Mapping

QTL mapping was performed on 960 haploid F1 segregants generated from a cross between BY and RM, which were a gift from Leonid Kruglyak (*24*). The segregants, arrayed in ten 96-well plates, were transferred in parallel with a disposable plastic pinner into 200 μL CSM pH 4.7 + 0.05% gelatin media either with K28 (10 plates) or without K28 (additional 10 plates), as described in the “Liquid Growth Assay for Killer Toxin Resistance” section above. Growth was measured in parallel over 63 hours on a SPECTROstar Omega microplate reader equipped with a plate stacker. The K28 resistance of each segregant was quantified as the ratio of AUC_+K28_/AUC_-K28_, as described above. QTL mapping was performed by calculating the logarithm-of-odds (LOD) score for each of 28,220 biallelic markers, defined as

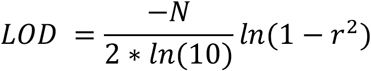

where *N* = 912 is the number of segregants after filtering out ill-behaved segregants, and *r* is the Pearson correlation between the K28 resistance phenotype and the genotype of each segregant at that biallelic marker; see page 454 of Lynch and Walsh (*44*). The genotypes are coded with -1 representing that the segregant had the RM allele and +1 signifying the BY allele, as determined by Bloom, et al. (*24*). The significance threshold was determined by a permutation test: QTL mapping was done 1000 times on permutations of the strain phenotypes relative to the strain genotypes, and the top LOD score was taken from each mapping. Among these LOD scores, the 950th highest value was taken as the 5% family-wise error rate (FWER) threshold for significance. Three QTLs were detected with peak LOD scores above this threshold. 95% confidence intervals for the location of the peak of these QTLs were determined by bootstrap with 1000 samplings with replacement (*45*), taking the peak marker per QTL-containing chromosome from each bootstrap mapping, ordering them from left to right, and determining the confidence interval as the span between the 2.5th and 97.5th percentile bootstrap peaks.

### Cloning of *KTD1* alleles

*KTD1* alleles tested in figures 2D, 2E, and 3B were amplified from genomic DNA using PfuUltraII Fusion HS DNA Polymerase (Agilent, Santa Clara, CA, USA) and primers listed in Table S8. Genomic DNA was isolated using DNeasy Blood & Tissue Kit (Qiagen) and following Qiagen’s Supplementary Protocol “Purification of total DNA from yeast using the DNeasy Blood & Tissue Kit”. Primers used for amplification are listed in Table S8. Amplicons were Gibson assembled (New England Biolabs, Ipswich, MA, USA) with pRS313 (MSp64) digested with XbaI-HF and XhoI-HF (New England Biolabs) or with MSp101 (*KTD1*_BY_) digested with NotI-HF and ApaI-HF (New England Biolabs).

### Chimeragenesis

All chimeras between *KTD1* and *UIP3*, as well as *KTD1* and *UIP3* themselves, were expressed from the *KTD1* promoter (nucleotide position −311 to −1 relative to the *KTD1* start codon) and terminator (position +1 to +230 relative to the *KTD1* ORF). Fragments were assembled via Gibson assembly. “Insert” PCR amplicons were generated using the high-fidelity PfuUltraII Fusion HS DNA Polymerase, while the longer “plasmid” PCR amplicons were generated using Herculase II Fusion DNA Polymerase (Agilent). Primer sequences used for chimeragenesis are listed in Table S8.

To construct chimeras, first, we used the *KTD1* plasmid, MSp101, to construct the *UIP3* plasmid, MSp160: we Gibson assembled the *UIP3* coding sequence amplified from BY genomic DNA with an amplicon of plasmid MSp101 to replace the *KTD1* coding sequence with *UIP3*. To generate chimeras, we amplified a “plasmid” fragment from MSp101 that contained a segment of *KTD1* along with the plasmid backbone. We amplified the corresponding “insert” fragment from MSp160 which contained the desired *UIP3* segment. The chimera-specific primers used to produce each plasmid-insert pair of PCR amplicons gave them approximately 20 bp of homology at each end. These fragments were Gibson assembled. Chimera transition points were selected to according to the conservation- and topology-based domains of DUP240 proteins defined by Poirey, et al. (*25*). Each plasmid was transformed into BY *ktd1Δ* and tested as described in the section “Killer Toxin Resistance Assay in Liquid Media.”

### Analysis of positive selection

Assemblies of the genomes of the 16-isolate panel generated by Moleculo technology (Illumina, San Diego, CA, USA) were obtained from Bloom, et al. (*17*). The 10 DUP240 genes from the reference *S. cerevisiae* genome were used as BLASTn (BLAST+ v2.10.0) queries against each of the assembled genomes using default search parameters. Hits were recursively added to the search query until no new hits were found, leading to a set of 15 new DUP240 orthologs, none of which had greater than 96% nucleotide identity to another DUP240. Sequences are available on GitHub and have been submitted to the Saccharomyces Genome Database.

A codon-based alignment of the DUP240 genes was generated using PRANK (v.170427) (*46*) with the gaprate parameter set to 0.000001 and a gap extension penalty of 0.001. As the codeML module of PAML does not estimate dN/dS values for codons aligned to gaps, DUP240 genes that created lots of gaps in the alignment were removed, resulting in a final alignment of 18 DUP240 orthologs (fig. S11). Analysis of BLASTn results showed that some of these orthologs are recombinants, whose inferred phylogeny may be incorrect. We performed recombination analysis using GARD (*47*) to predict recombination breakpoints. For breakpoints that occurred in the middle of a codon, we rounded to the nearest codon boundary. Seven breakpoints were identified, which we used to break the alignment into 8 segments for further analysis.

Phylogenetic trees were generated on each segment with FastTree (2.1) (*48*) with *-gtr -gamma* options. We then used the codeml module in PAML (v 4.9j) to generate models M7 (no positive selection) and M8 (with positive selection) on each segment with the corresponding segment-specific alignment and phylogenetic tree. The two models were compared by computing the likelihood-ratio test statistic,

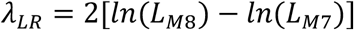

which is chi squared-distributed with two degrees of freedom. The results are summarized in Table S5.

### Toxin Adsorption

To test the ability of K28 toxin to bind to the surface of yeast cells, we adapted the toxin-cell binding assay described in Carroll, et al. and Breinig, et al. (*28, 29*). Strains MSY94, 96-100, and 104 (*n* = 4 biological replicates) were grown to saturation for 48 h in 30 mL CSM-His (pH 7.0) at 30 C, shaking at 200 RPM. 200 mL of toxic and non-toxic supernatants were prepared as described earlier, with 24 h incubation time. Strain MSY26 (RM) was grown for 24 h in YPD. 10^9^ cells (50 OD_600_ units) of each experimental strain were pelleted at 3000 RPM. Supernatant was mostly aspirated, leaving approximately 0.5-0.7 mL media in the tube. Pellet was resuspended in the remaining supernatant and placed on ice.

5 mL of toxic supernatant was added to each cell suspension and incubated for 15 min on a nutator at approximately 1 RPM. As controls, we also incubated cell-free toxic and non-toxic supernatants. Suspensions and cell-free controls were then filtered using syringes with 0.22 µm Nalgene syringe filters (Thermo Fisher Scientific), generating filtrates of dissolved toxin that had not adsorbed to cell surfaces. The toxin adsorption and filtration steps were done at 4 C. The degree of toxin adsorption was quantified as the area under the growth curve (AUC_+K28_) of the K28-sensitive strain RM in 160 μL filtrate + 40 μL 5xCSM pH 4.7 for 48 h (fig. S7).

### Software, Statistical Analysis, Data Deposition, and Data Visualization

Statistical analysis and data visualization were performed using custom R scripts, available at https://github.com/ilya-andreev2/killer-virus. For experiments with comparisons among many strains, statistical analysis was performed using one-way ANOVA followed by Tukey’s post-hoc HSD test. For experiments comparing specific pairs of strains, statistical analysis was performed with Welch’s two-sample *t*-tests.

Growth curves were generated using the *geom_smooth* function in the R package ggplot2 (version 3.3.5) (*49*). Area under the curve (AUC) was calculated using the *AUC* function (trapezoid method) from the R package DescTools v.0.99.42. Neighbor-joining phylogenetic trees of 1,011 sequenced yeast strains (*30*) were generated using the function *bionj* from the R package ape (version 5.3) (*50*). The maximum edge length was set to 0.7 to truncate a highly diverged clade of yeast strains. Jittered 1-dimensional scatter plots were generated using the *ggboxplot* function in R package ggpubr. Protein sequence alignments were generated in Clustal Omega (*51*). The structure of Ktd1p was predicted using AlphaFold (version 2.0.0) (*37*) and visualized in PyMOL (version 2.3.0 Open-Source), with the position of the transmembrane helices shown according to Poirey, et al. (*25*).

Moleculo-based genome assemblies of the strains in our 16-isolate panel are being submitted to the NCBI genome database, and non-reference DUP240 genes have been submitted to the Saccharomyces genome database (SGD) named *DFP11* through *DFP25* (*D*UP240 *f*amily *p*rotein).

---

**Figure S1.**
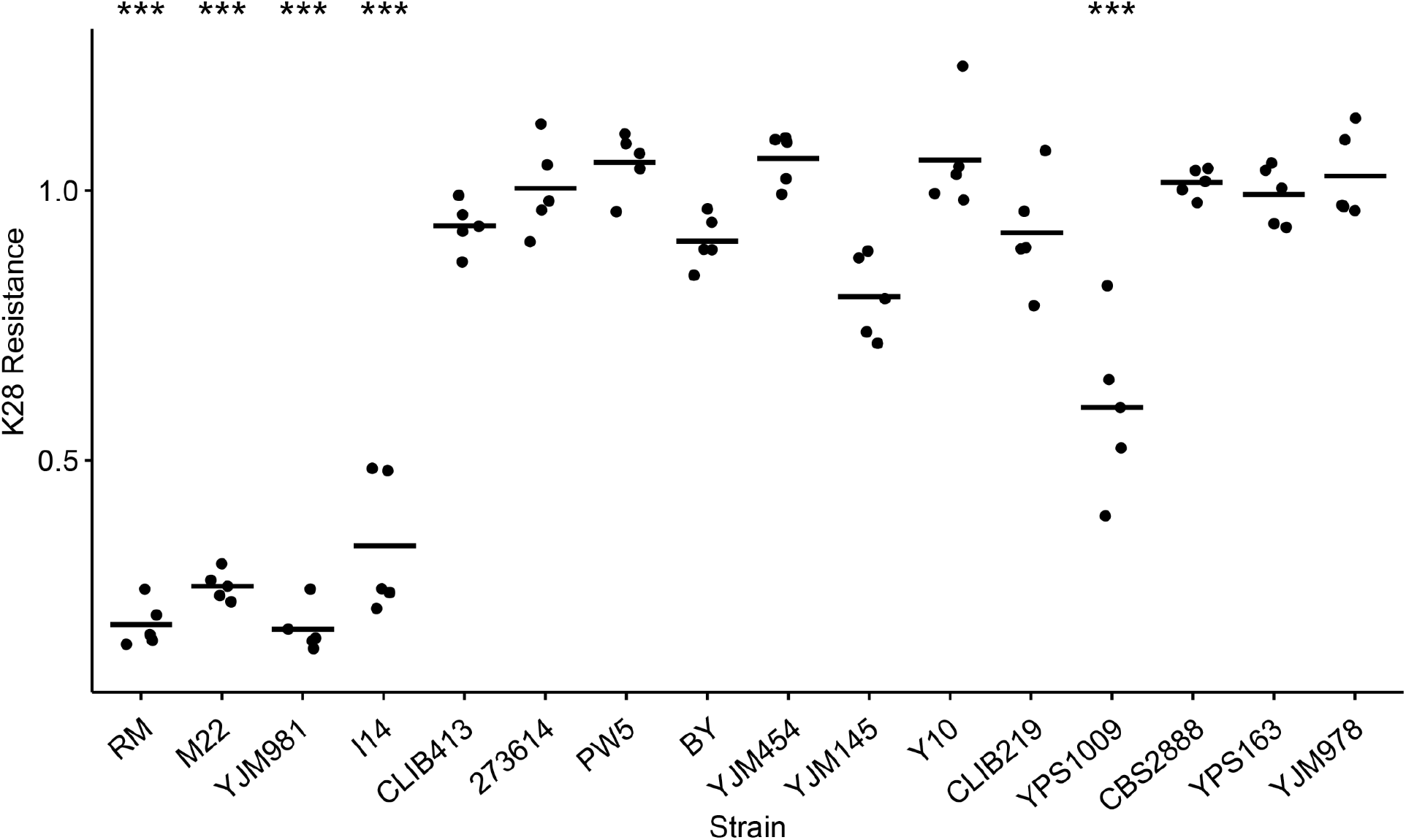
Statistical analysis of K28 resistance of 16 diverse S. cerevisiae strains. From the growth curves in figure 1B, K28 resistance was quantified as AUC_+K28_ / AUC_-K28_ with *n* = 5 biological replicates per strain. One-way ANOVA indicated a statistically significant effect of strain on K28 resistance (K28 resistance ∼ strain). Tukey’s post-hoc HSD test was performed to test for the difference in mean K28 resistance between all pairs of strains. Strains with significantly lower K28 resistance are marked with asterisks (\*\*\**P* < 6.6×10^-6^ in comparison to BY). Horizontal bars indicate the sample means.

**Figure S2.**
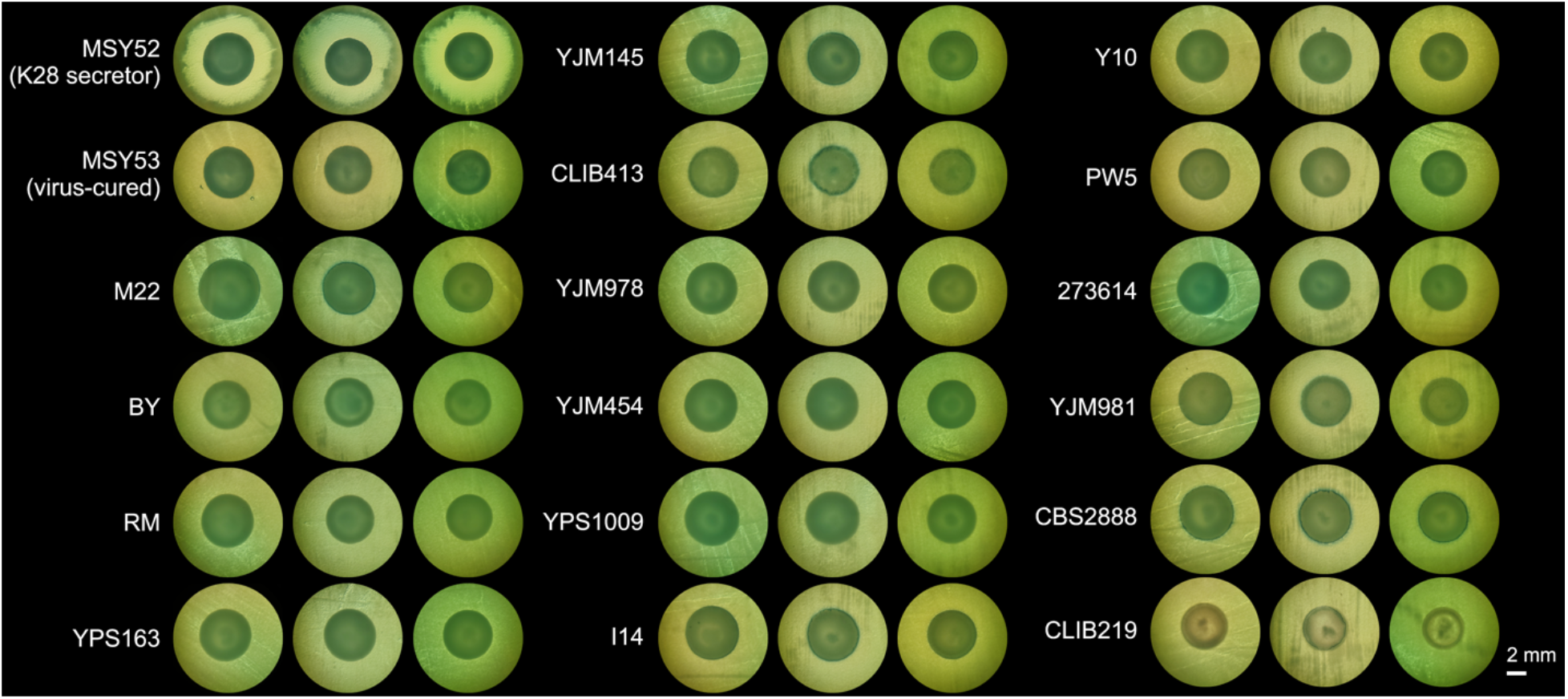
Phenotypic Virus Typing – Halo Assay. The 16-isolate panel was tested for production of K28 toxin by whether they killed the hypersensitive yeast strain 192.2d. Triplicate spots of each strain were grown on a lawn of 192.2d cells; strains producing K28 will generate a halo of cleared 192.2d cells (*28*). MSY52, a diploid *ski2**Δ*** strain infected by M28, was spotted as a positive control; MSY53, a virus-cured derivative of MSY52, was spotted as a negative control.

**Figure S3.**
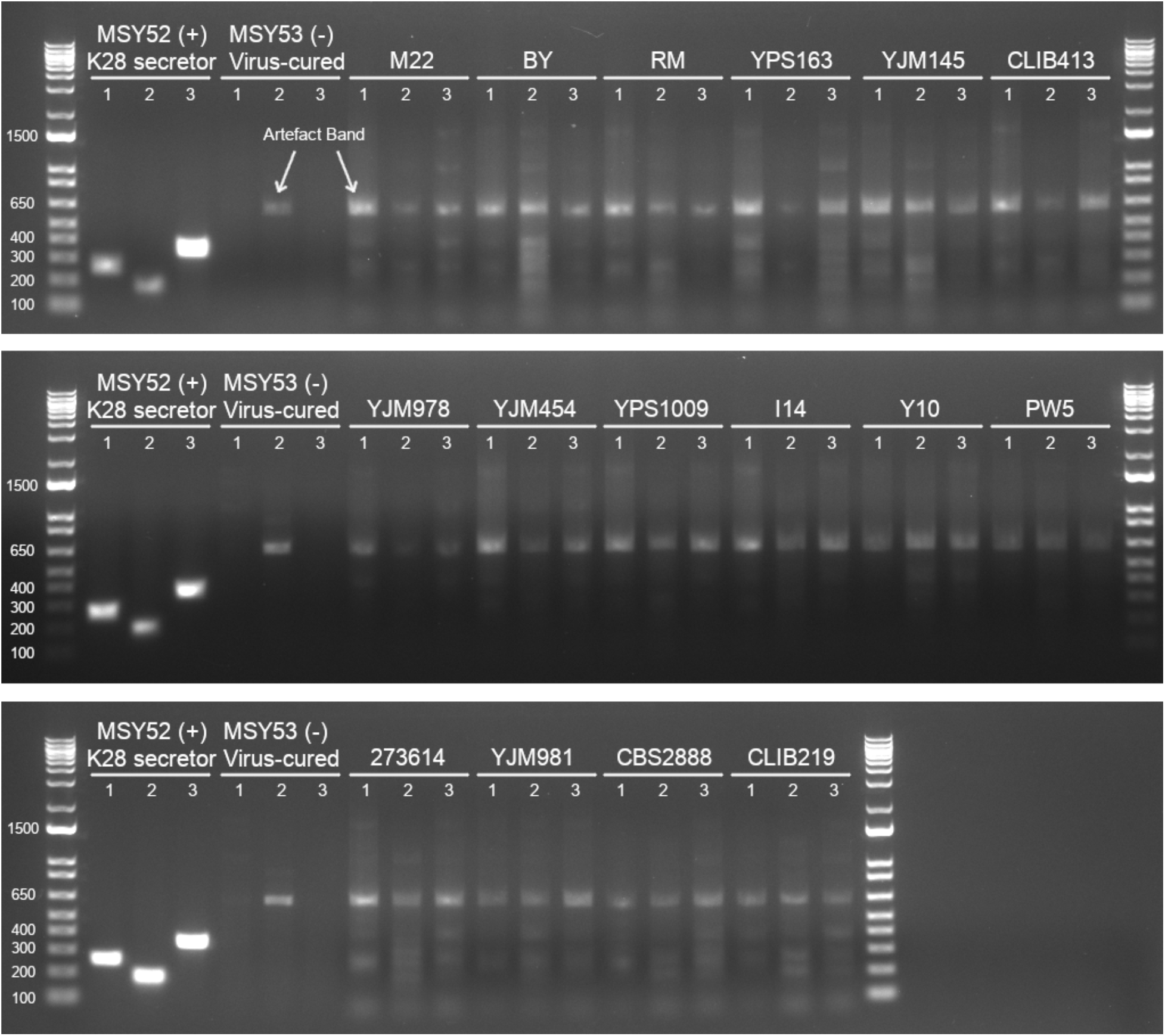
RT-PCR-based Virus Typing. Three separate PCRs were performed on cDNA prepared from the 16-isolate panel and two control strains. Each PCR was designed to amplify a part of the M28 virus’s genome sequence (*18*). Primer sequences are provided in Table S8 (PCR 1: expected size of 253 bp; PCR 2: expected size of 180 bp; PCR 3: expected size of 350 bp). Ladder is 1kb Plus (Invitrogen). Shown gel is representative of *n* = 3 biological replicates, derived from three colonies of each strain.

**Figure S4.**
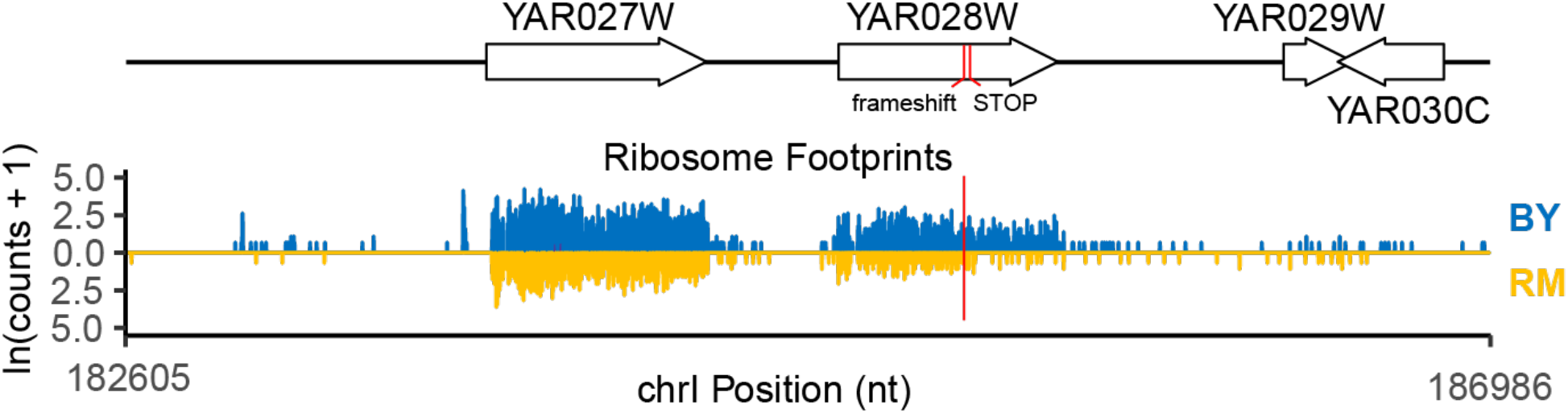
Ribosome profiling of BY and RM strains near the YAR027W (*UIP3)* and YAR028W (*KTD1*) genes. Log-transformed counts of ribosome footprints, as determined by Albert, et al. (*27*), are shown in a window of chromosome I encompassing *UIP3* (YAR027W) and *KTD1* (YAR028W). Footprints are plotted in blue for BY and gold for RM. The positions of deleterious mutations in RM are marked with a vertical red line.

**Figure S5.**
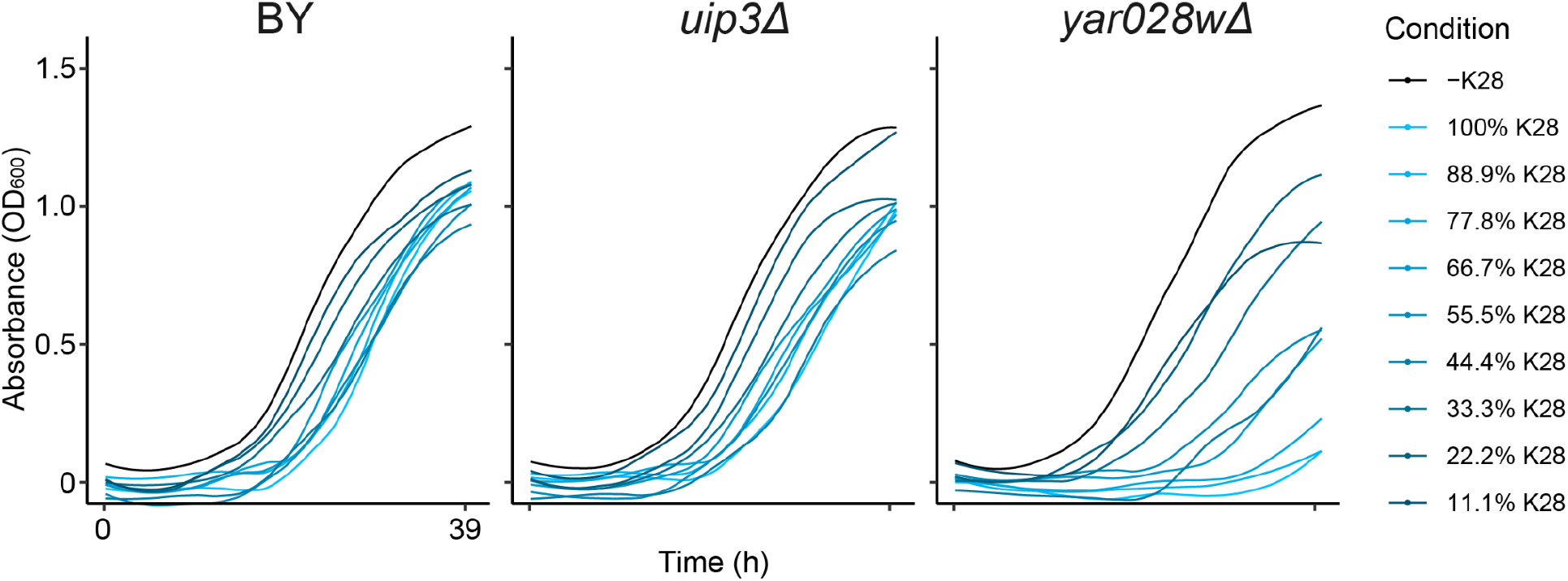
Deletion of *UIP3* had no effect on K28 resistance. BY, BY *uip3Δ*, and BY *yar028wΔ* (*ktd1Δ*) were grown in media containing different concentrations of K28, generated by mixing the supernatants of K28-producing (MSY52) and virus-free (MSY53) yeast strains. Legend indicates the percentage of toxic supernatant (X%) diluted in non-toxic supernatant (100% - X%).

**Figure S6.**
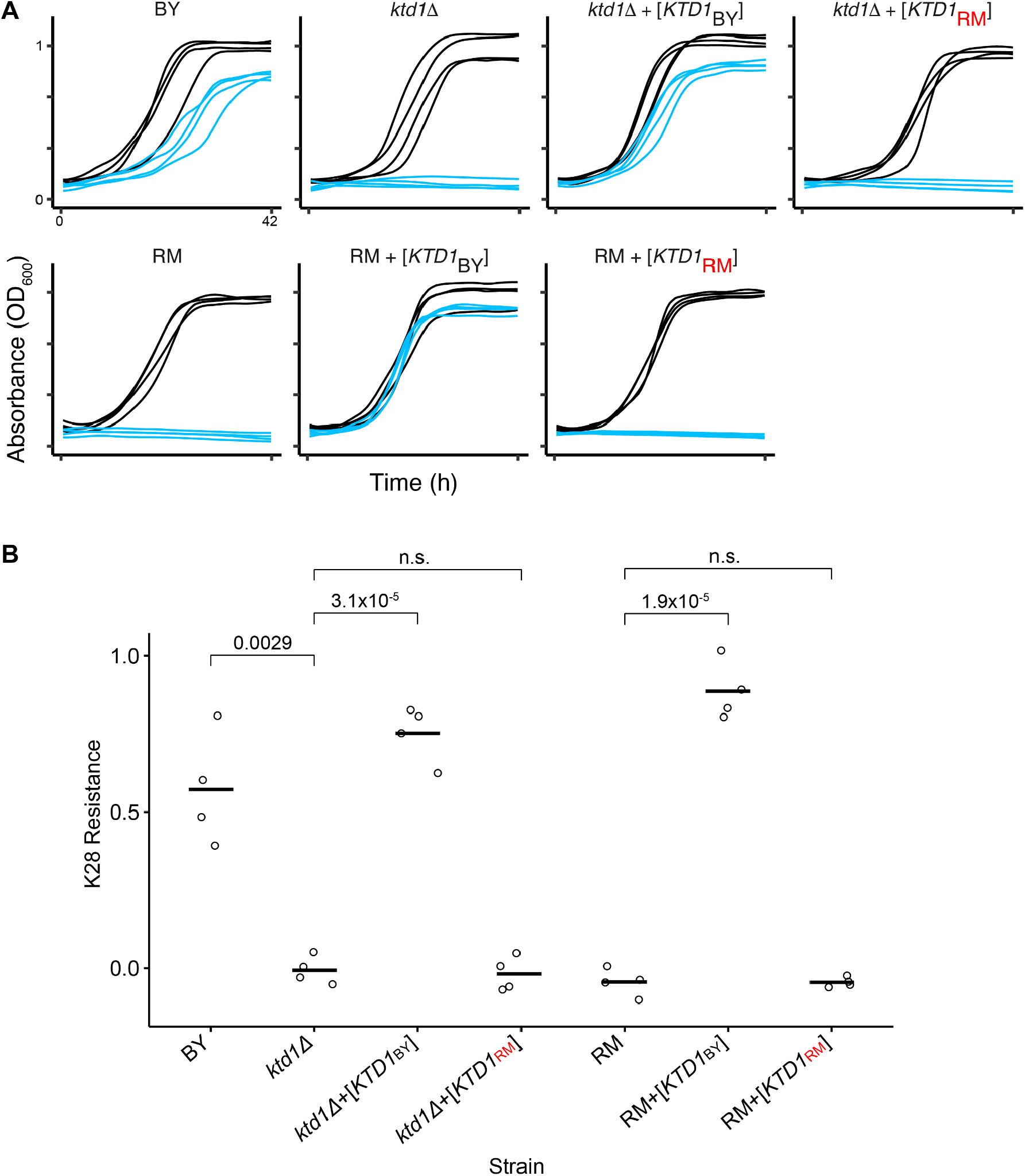
Expression of the RM allele of *KTD1* did not protect against K28. **(A)** Growth curves of BY, RM, and BY *ktd1Δ* are shown, expressing either the BY or RM allele of *KTD1*, or empty vector. Blue curves correspond to growth in media containing K28; black curves correspond to growth in media without K28, with *n* = 4 biological replicates per strain. A subset of these curves are also shown in figure 2D and E. **(B)** From the growth curves shown in part A, K28 resistance was quantified as AUC_+K28_ / AUC_-K28_. Welch’s two-sample *t*-tests were used to compare levels of K28 resistance between strains.

**Figure S7.**
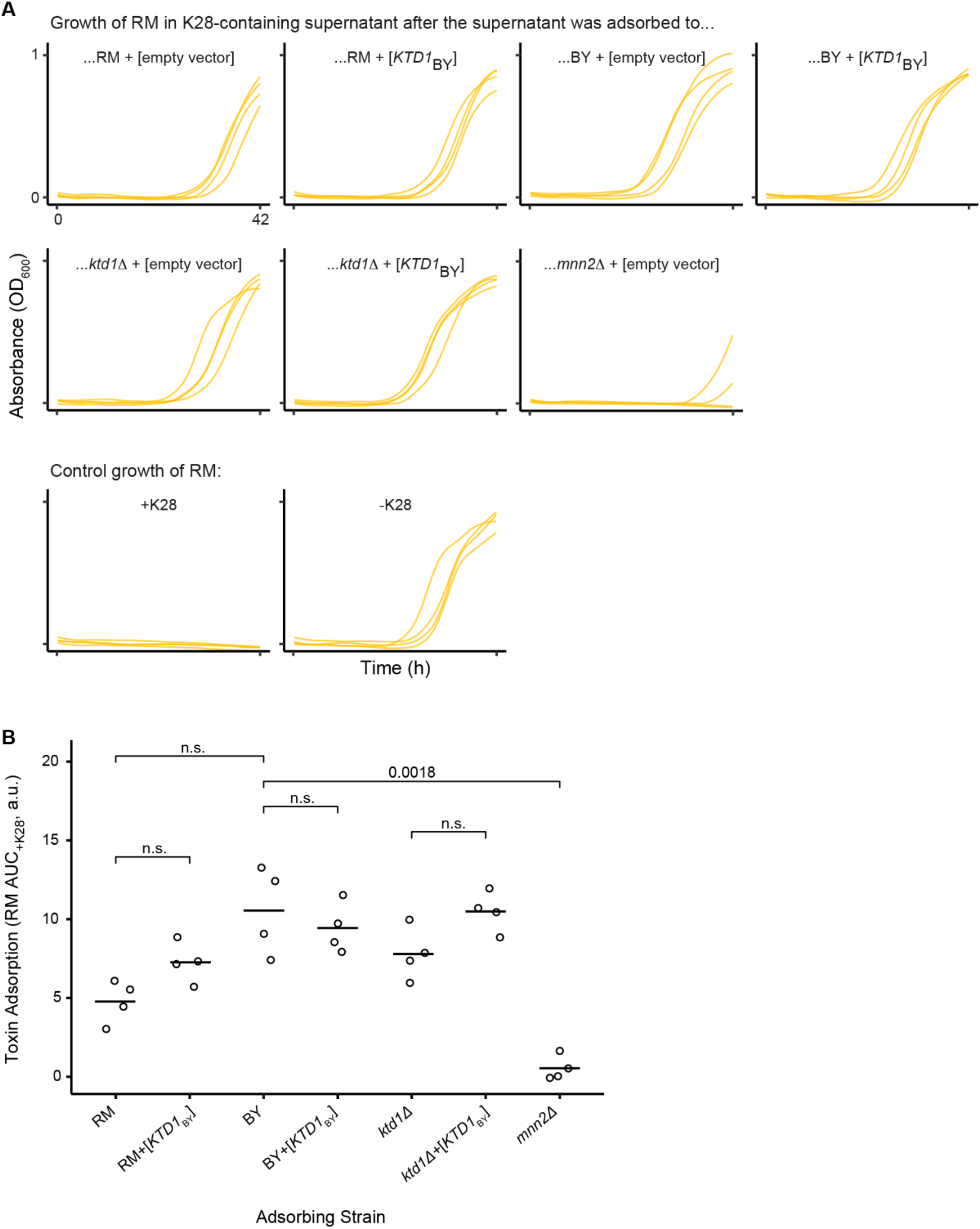
Effect of Ktd1p on adsorption of K28 to the cell surface. **(A)** Supernatant from K28-secreting MSY52 was applied to cells of the genotypes shown (*n* = 4 biological replicates) for 15 minutes, after which the supernatant was filter-sterilized to remove the cells and any K28 adsorbed onto their surface. The level of remaining K28 toxin was determined by the ability of the K28-sensitive RM strain to grow in the supernatant. As controls, the growth of RM in supernatant of MSY52 (with toxin) and MSY53 (toxin-free) without an adsorption step is shown. **(B)** Quantification of the inhibition of RM’s growth by non-adsorbed K28, measured as AUC_+K28_ (vertical axis) from the growth curves in part A. Higher values indicate more toxin was adsorbed, hence the growth of RM was less inhibited. Statistical analyses were performed using Welch’s two-sample one-tailed *t*-tests, with the alternative hypothesis that *KTD1*_BY_ leads to decreased toxin adsorption.

**Figure S8.**
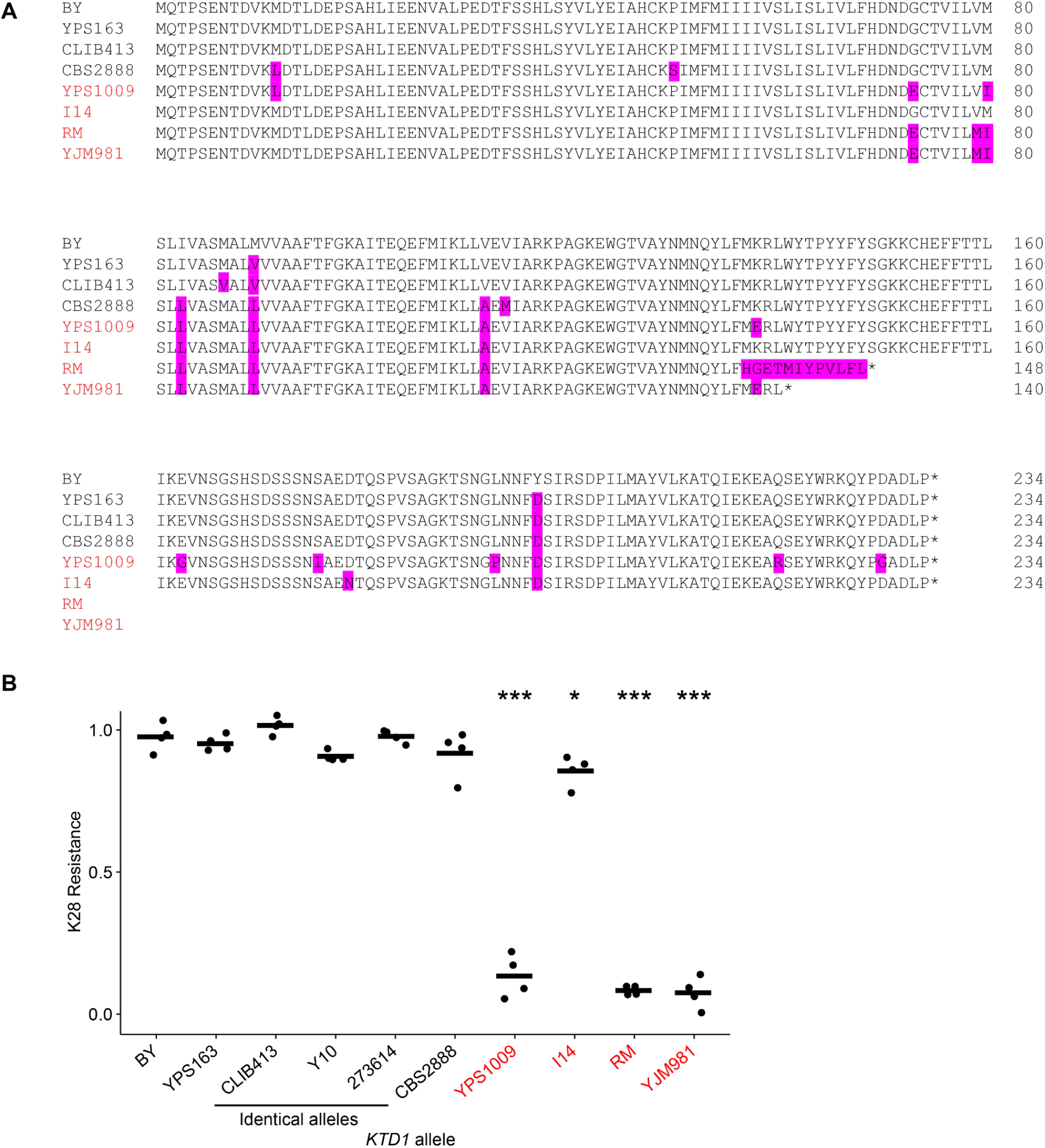
*KTD1* alleles from the 16-isolate panel. Strain names are colored according to K28 resistance as determined in Fig. 1B (black, resistant; red, sensitive). **(A)** Amino acids that differ from the reference sequence (BY) are highlighted in magenta. Strains Y10 and 273614 had *KTD1* alleles identical to CLIB413 and are not shown here. Sequences were determined by Illumina Moleculo sequencing performed in Bloom, et al. (*17*). **(B)** From the growth curves shown in figure 3B of BY *ktd1Δ* expressing various *KTD1* alleles, K28 resistance was quantified as AUC_+K28_ / AUC_-K28_ with *n* = 4 biological replicates per strain. ANOVA followed by Tukey’s HSD identified the six alleles from resistant strains and *KTD1*_I14_ as conferring more resistance than *KTD1*_YPS1009_, *KTD1*_RM_, and *KTD1*_YJM981_ (\*\*\**P* < 0.001 for all 21 pairwise comparisons). *KTD1*_I14_ conferred less resistance than *KTD1*_BY_ (\**P* < 0.05), *KTD1*_273614_ (*P* < 0.05), and *KTD1*_CLIB413_ (*P* < 0.01) at more moderate thresholds for statistical significance, but was not significantly different from the alleles from other resistant strains.

**Figure S9.**
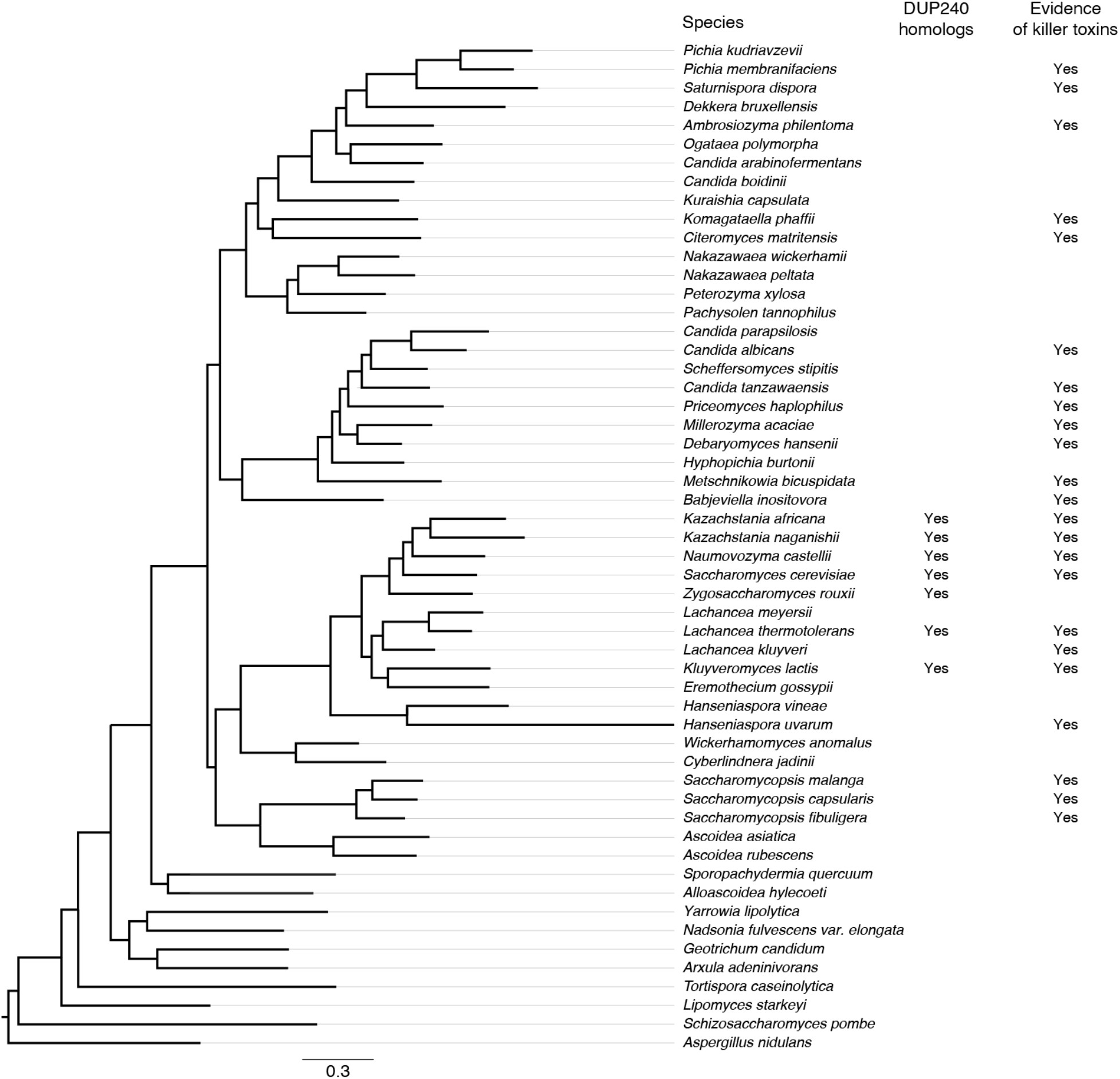
Species distribution of homologs of DUP240 genes. Homologs of DUP240 genes were found by performing tblastn searches against each genome shown on the tree, with the Ktd1p protein sequence as query. All negative hits were negative both with the default tblastn search parameters and with the tblastn Expect threshold relaxed to 0.5. Evidence of killer toxins is largely as reported by Krassowski, et al. (*52*), with the exception of *P. membranifaciens, K. africana, N. castellii,* and *H. uvarum* (*20, 53, 54*). Phylogenetic species tree modified with permission from Krassowski, et al.

**Figure S10.**
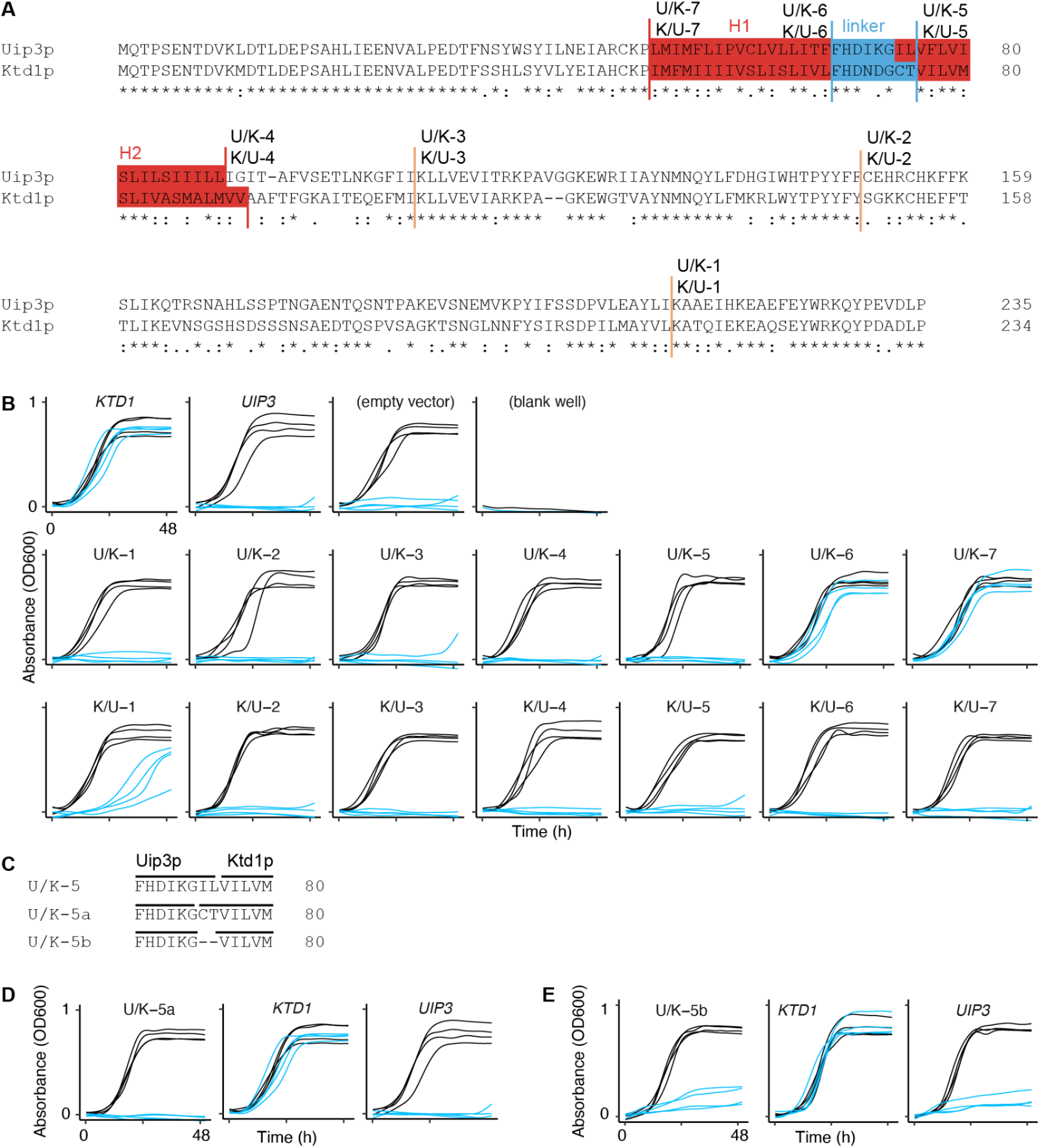
Analysis of chimeras between Uip3p and Ktd1p. **(A)** Alignment of Ktd1p and Uip3p. Chimera transition points are marked with vertical lines and the names of the chimeras transitioning there. The transmembrane helices H1 and H2 are colored in red and the linker between them is colored in blue. We used the positions of the transmembrane domains as defined in Poirey et al. (*25*). **(B)** The K28 resistance shown in figure 4A was quantified as AUC_+K28_ / AUC_-K28_ calculated from the growth curves shown here of BY *ktd1Δ* expressing the labeled chimeras, with *n* = 4 biological replicates per strain. Blue growth curves are of strains grown in media containing K28, and black growth curves are of strains grown in media lacking K28. **(C)** We additionally tested two alternative Uip3p-Ktd1p chimera transition points near the transition point used in U/K-5. U/K-5a transitions at a conserved glycine within the interhelix linker, and U/K-5b transitions from the end of Uip3p’s linker to the start of Ktd1p’s H2 transmembrane domain. **(D)** Growth curves of BY *ktd1Δ* expressing U/K-5a in media with and without K28. BY *ktd1Δ* expressing either *KTD1* or *UIP3* are shown for comparison. **(E)** Growth curves of BY *ktd1Δ* expressing U/K-5b in media with and without K28. BY *ktd1Δ* expressing either *KTD1* or *UIP3* are shown for comparison. In D and E, U/K-5a and U/K-5b conferred significantly less resistance than *KTD1* (*P* < 0.001 by Welch’s two-sample *t*-test) and were not distinguishable from *UIP3*. U/K-5a was tested in the same experiment as the chimeras shown in B; U/K-5b was tested separately.

**Figure S11.**
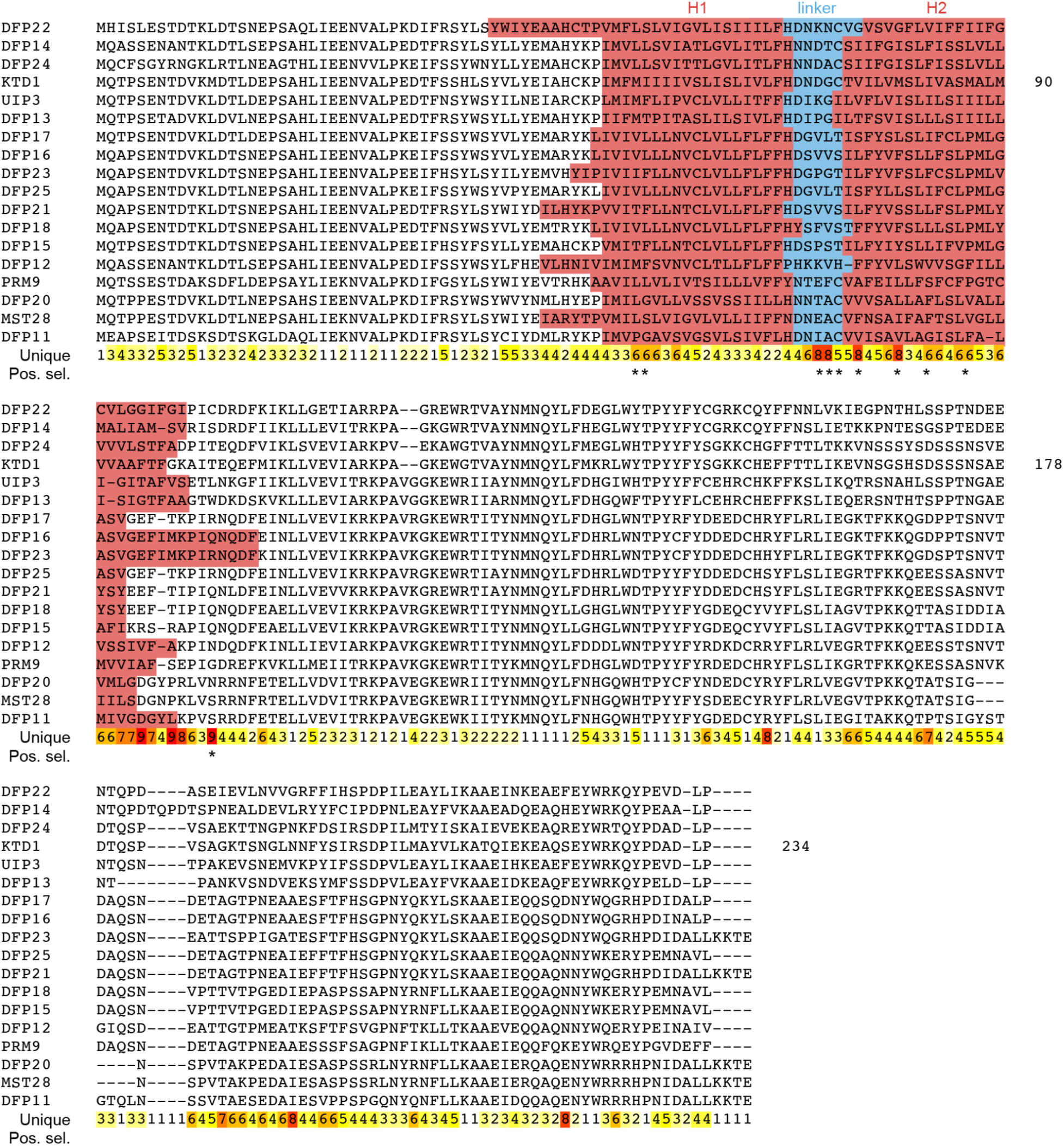
Regions of divergence between DUP240 proteins. In our 16-isolate panel we found 25 DUP240 genes; of these, the 18 full-length DUP240 proteins most similar to Ktd1p are shown in this alignment. DUP240 genes with no previous standard name were named DFP (*<u>D</u>*UP240 *f*amily *<u>p</u>*rotein). The locations of transmembrane domains TM1 and TM2 were determined by PHOBIUS (*31*). Along the bottom of the alignment we show the number of unique amino acids represented at each position in the alignment (“Unique”) and the sites identified as experiencing positive selection (“Pos. sel.”); see figure 4B.

**Table S1.** Origin of the S. cerevisiae strains in the 16-isolate panel surveyed in this study. Strains and their origin information from Bloom, et al., 2019 and Peter, et al., 2018 (*17, 30*). NA = information not available.

| Strain Name | Environmental / Geographical Origin |
| --- | --- |
| RM11-1a (MATalpha) | Zinfandel vineyard / CA, USA |
| M22 (MATalpha) | Vineyard / Italy |
| YJM981 (MATalpha) | Vagina / Italy |
| I14_1b (MATa) | Vineyard soil / Italy |
| CLIB413_1b (MATa) | Fermenting rice / China |
| 273614N (MATa) | NA / NA |
| PW5_b (MATalpha) | Raphia palm wine / NA |
| BY (MATa) | (Prototrophic version of a common lab strain) |
| YJM454 (MATa) | Human, clinical / NA |
| YJM145 (MATalpha) | AIDS patient / NA |
| Y10 (MATalpha) | Coconut / Philippines |
| CLIB219 (MATalpha) | Wine / Russia |
| YPS1009 (MATalpha) | Exudate, *Quercus* sp. (oak) / NA |
| CBS2888 (MATa) | Soil / South Africa |
| YPS163 (MATa) | Soil beneath *Quercus rubra* (oak) / PA, USA |
| YJM978 (MATalpha) | NA / NA |

**Table S2.** QTLs identified from linkage mapping of K28 resistance variation in a BY × RM cross. 912 segregants from a BY × RM cross were phenotyped for resistance to K28 killer toxin, measured as AUC_+K28_/AUC_-K28_. The genotypes at 28,220 biallelic markers spanning the genome were associated with K28 resistance, generating the LOD plot in figure 2B. We identified three QTLs that passed a 5% family-wise error rate (FWER) LOD threshold of 3.55, computed from 1000 iterations of randomly assigning phenotypes to segregants and calculating the top LOD score. The left and right markers denote 95% confidence intervals for the position of the LOD peak marker. These were determined from the positions of peak LOD scores on chromosomes I, XII, and XIV from 1000 bootstrap samplings.

| QTL | Chr | Peak Marker Position | Left Marker | Right Marker | LOD Score |
| --- | --- | --- | --- | --- | --- |
| 1 | I | 184648 | 180646 | 184686 | 49.72978 |
| 2 | XII | 745464 | 250697 | 967302 | 4.11199 |
| 3 | XIV | 468488 | 467028 | 485549 | 11.36227 |

**Table S3.**
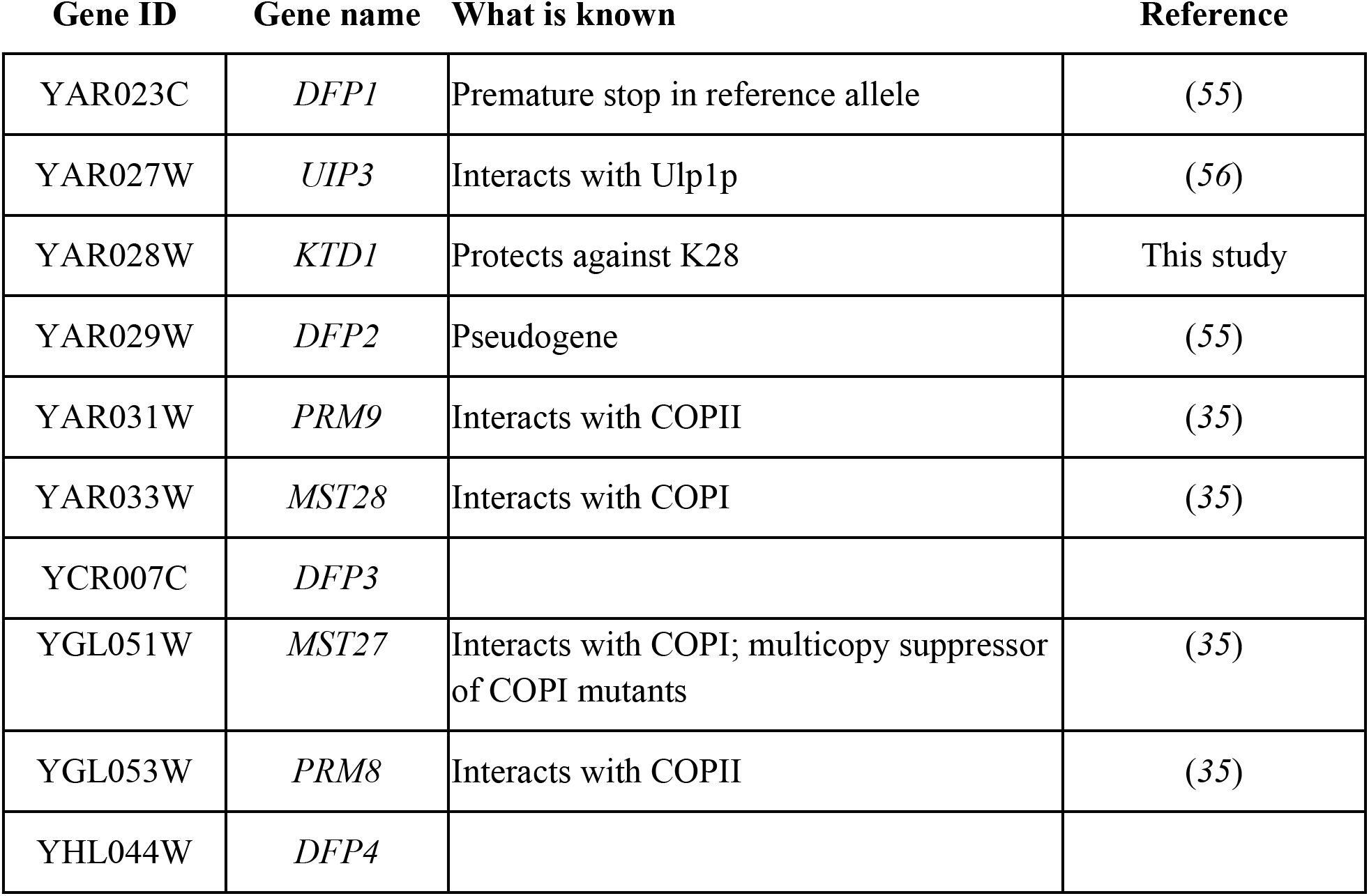
DUP240 genes in the yeast reference genome. We have named the four remaining unnamed DUP240 genes in the *S. cerevisiae* reference genome as *DFP* (*<u>D</u>*UP240 *f*amily *<u>p</u>*roteins).

| Gene ID | Gene name | What is known | Reference |
| --- | --- | --- | --- |
| YAR023C | *DFP1* | Premature stop in reference allele | (55) |
| YAR027W | *UIP3* | Interacts with Ulp1p | (56) |
| YAR028W | *KTD1* | Protects against K28 | This study |
| YAR029W | *DFP2* | Pseudogene | (55) |
| YAR031W | *PRM9* | Interacts with COPII | (35) |
| YAR033W | *MST28* | Interacts with COPI | (35) |
| YCR007C | *DFP3* |  |  |
| YGL051W | *MST27* | Interacts with COPI; multicopy suppressor of COPI mutants | (35) |
| YGL053W | *PRM8* | Interacts with COPII | (35) |
| YHL044W | *DFP4* |  |  |

**Table S4.**
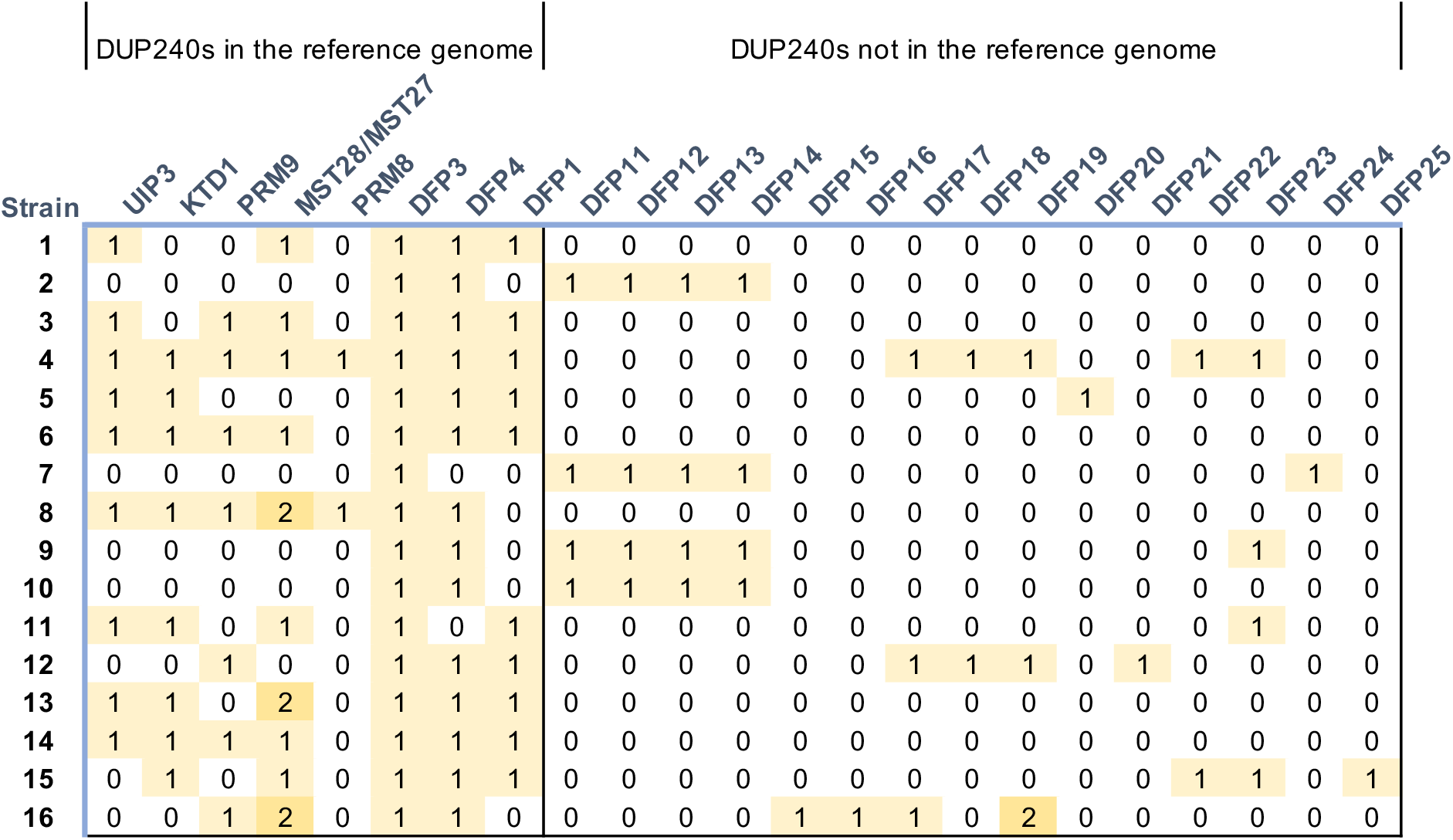
Presence and absence of DUP240 homologs in the genomes of strains in the 16-isolate panel. BLASTn was used to find all sequences homologous to any of the 10 DUP240 genes from the reference genome. Sequences with greater than 96% nucleotide identity were grouped as alleles; those with less than 96% nucleotide identity were called distinct genes. The sequences of 15 non-reference DUP240 genes have been deposited at SGD.

**Table S5.** Analysis of rapidly evolving sites in the DUP240 family. Sites under positive selection were identified using the codeML module in PAML. The DUP240 ortholog alignment (shown in fig. S11) was divided into segments according to results of DNA recombination breakpoint analysis using GARD. Model M7 (negative or neutral selection) and Model M8 (negative, neutral, or positive selection) were compared using the likelihood-ratio test, followed by the chi-squared significance test to determine where Model M8 should be accepted, which would indicate the presence of a signature of positive selection in the region. Segments 48-89 and 100-124 were identified as having such signatures. Bayes Empirical Bayes (BEB) analysis on these segments revealed sites under positive selection (dN/dS > 1) with posterior probability Pr(dN/dS > 1) greater than 0.95. Codon positions are with respect to *KTD1*.

| Segment<br>(*KTD1*<br>a.a.) | Mean dN/dS<br>(Site model) | Likelihood-ratio<br>test statistic<br>$2*[\ln(L_{M8})-\ln(L_{M7})]$ | *p*-value of<br>chi<sup>2</sup> test<br>(df = 2) | *KTD1* sites under positive selection<br>with Pr(dN/dS>1) using BEB<br>(* > 0.95, ** > 0.99) |
| --- | --- | --- | --- | --- |
| 1-18 | 0.8200 (M7) | 2.69 | 0.261 | N/A |
| 19-36 | 0.2911 (M7) | 0.90 | 0.637 | N/A |
| 37-47 | 0.4695 (M7) | 0.62 | 0.733 | N/A |
| 48-89 | 2.0006 (M8) | 40.28 | **1.795×10<sup>-9</sup>** | 55I, 0.986 (*)<br>56I, 0.991 (**)<br>72D, 1.000 (**)<br>73G, 0.988 (*)<br>74C, 0.973 (*)<br>76V, 0.997 (**)<br>80M, 0.992 (**)<br>83I, 0.999 (**)<br>87M, 0.970 (*) |
| 90-99 | 1.0000 (M7) | 3.61 | 0.165 | N/A |
| 100-124 | 0.6415 (M8) | 7.66 | **0.021** | 102T, 0.998 (**) |
| 125-210 | 0.5975 (M7) | 2.45 | 0.294 | N/A |
| 211-234 | 0.4283 (M7) | 0.17 | 0.919 | N/A |

Tables included as separate files:

**Table S6.** Strains used in this study.

**Table S7.** Plasmids used in this study.

**Table S8.** Oligonucleotides used in this study.

